# Meta-Signer: Metagenomic Signature Identifier based on Rank Aggregation of Features

**DOI:** 10.1101/2020.05.09.085993

**Authors:** Derek Reiman, Ahmed A. Metwally, Jun Sun, Yang Dai

## Abstract

**Background:** The advance of metagenomic studies provides the opportunity to identify microbial taxa that are associated to human diseases. Multiple methods exist for the association analysis. However, the results could be inconsistent, presenting challenges in interpreting the host-microbiome interactions. To address this issue, we introduce Meta-Signer, a novel <u>Meta</u>genomic <u>Sign</u>ature Identifi<u>er</u> tool based on rank aggregation of features identified from multiple machine learning models including Random Forest, Support Vector Machines, LASSO, Multi-Layer Perceptron Neural Networks, and our recently developed Convolutional Neural Network framework (PopPhy-CNN). Meta-Signer generates ranked taxa lists by training individual machine learning models over multiple training partitions and aggregates them into a single ranked list by an optimization procedure to represent the most informative and robust microbial features. Meta-Signer can rank taxa using two input forms of the data: the relative abundances of the original taxa and taxa from the populated taxonomic trees generated from the original taxa. The latter form allows the evaluation of the association of microbial features at different taxonomic levels to the disease, which is attributed to our novel model of PopPhy-CNN.

**Results:** We evaluate Mega-Signer on five different human gut-microbiome datasets. We demonstrate that the features derived from Meta-Signer were more informative compared to those obtained from other available feature ranking methods. The highly ranked features are strongly supported by published literature.

**Conclusion:** Meta-Signer is capable of deriving a robust set of microbial features at multiple taxonomic levels for the prediction of host phenotype. Meta-Signer is user-friendly and customizable, allowing users to explore their datasets quickly and efficiently.

## Background

Recent metagenomic studies of the gut microbiome have linked dysbiosis to many human diseases [1, 2, 3]. A metagenomic sample is typically represented by its microbial taxonomic composition using microbial taxa at one of the taxonomic levels, i.e., Super-kingdom, Phylum, Class, Order, Family, Genus, and Species. The identification of microbial taxa associated with the human disease has been one of important efforts in metagenomics data analysis [4]. Procedures used in various metagenomic studies use parametric or non-parametric statistical tests to detect differentially abundant individual taxa between disease and control groups [5, 6, 7, 8, 9]. These type of methods can potentially miss taxa with weak associations which can together present strong statistical association. In order to capture group association, several methods are proposed by exploring related taxa on a phylogenetic taxonomic tree. For example, a concept of variable fusion was introduced to bring two closely related taxa on the tree into a Lasso linear regression model [10]. OMiAT, a statistical framework, combines tests of all upper- and lower-level taxa to generate a microbiome comprehensive association mapping (MiCAM) [11]. A two-stage microbial association mapping framework (massMap) was proposed to perform individual test with the taxonomic groups detected by their novel microbial group test procedure [12]. massMap was shown to be statistically more powerful than two other aggregate-based methods, e.g., the Microbiome Regression-Based Kernel Association Test (MiRKAT) [13] and another sophisticated association test based on the generalized taxon proportion (MiSPU) [14]. These elaborated statistical testing methods enhance the detection of the microbial group association. However, they may still fail to detect complex multivariate non-linear associations. On the other hand, there are also concerns that introducing interactions between taxa may result in spurious associations, which can be removed by a procedure based on maximum entropy models [15].

Alternative approaches of using machine learning (ML) models have been advocated for the prediction of the host phenotype [16, 17]. This is motivated by the findings that a microbial signature for the host phenotype may be complex, involving simultaneous over- and under-representations of multiple microbial taxa potentially interacting with each other. Classical ML models, such as Random Forest (RF), Least Absolute Shrinkage and Selection Operator (LASSO) and other regression-based models with feature selection, Support Vector Machines (SVMs), and ensemble-based models have been applied [18, 10, 19, 17, 20]. The advanced use of a phylogenetic or taxonomic tree to imprint relevant biological knowledge in deriving meaningful features for metagenomic data has been seen in multiple ML models [21, 22]. These studies have shown that the traditional ML models using the hierarchical structure of the taxonomic tree can lead to better classification performance over the use of only original taxonomic features, further supporting the idea of generating microbial signature of taxa at distinct taxonomic levels.

In the effort of exploring high-order interactions between microbial taxa to benefit host phenotype prediction, deep neural networks (DNNs) have been proposed due to their ability of exploring complex relationship among features. Multilayer perceptron neural network (MLPNN) using relative taxa abundance vectors [23] was shown to outperform other DNNs and have similar predictive power as RF. More recently, a convolutional neural network (CNN) architecture was proposed to explore the distance between nodes on a phylogenetic tree by the patristic distance (the sum of the lengths of all branches connecting two OTUs on the tree) [24]. Their approach, i.e., Ph-CNN, was reported to outperform linear SVMs, RF and a baseline fully connected MLPNN on synthetic data using gut metagenomic data from 222 inflammatory bowel disease (IBD) patients and 38 healthy subjects. Another CNN-based framework (PopPhy-CNN) was developed by our group by designing an input format of a 2D matrix representing the taxonomic tree populated with the relative abundance of microbial taxa in a metagenomic sample [25, 26]. This conversion empowers CNNs to explore the spatial relationship of the taxonomic annotations on the tree and their quantitative characteristics in metagenomic data. PopPhy-CNN was shown competitive using nine metagenomic datasets of moderate size including the IBD dataset [26].

Despite of the promising results from the numerous ML and DNN approaches, a critical analysis tool, MetaPheno, showed that the predictive performance gains of recent models were small and that more focus should be placed on feature extraction [20]. In addition, to best of our knowledge, there is no reliable method that can systematically generate a robust set of features from a set of individual ML models.

In this work, we introduce a novel tool, Meta-Signer, <u>Meta</u>genomic <u>Sign</u>ature Identifi<u>er</u> based on rank aggregation of informative taxa learned from individual ML models. Meta-Signer uses RF, SVM, LASSO, MLPNN, and PopPhy-CNN models to evaluate importance of each microbial taxa and generates a ranked list of features per model. It aggregates all the ranked lists using a procedure based on the CrossEntropy method or the Genetic Algorithm [27]. Meta-Signer can extract features from two different ways: the original relative taxa abundance data and the taxa from taxonomic trees constructed from the original taxa. By providing a ranked list of the original taxa and a ranked list of the taxa from the taxonomic trees, the user receives a more robust overview of how the microbial community members are associated with the phenotype. Meta-Signer is user-friendly and easy to run. It provides a readable summary as an HTML output. Meta-Signer is distributed as a Python tool and available at https://github.com/YDaiLab/Meta-Signer.

## Implementation

The overview of the workflow is shown in Fig.1

**Figure 1.**
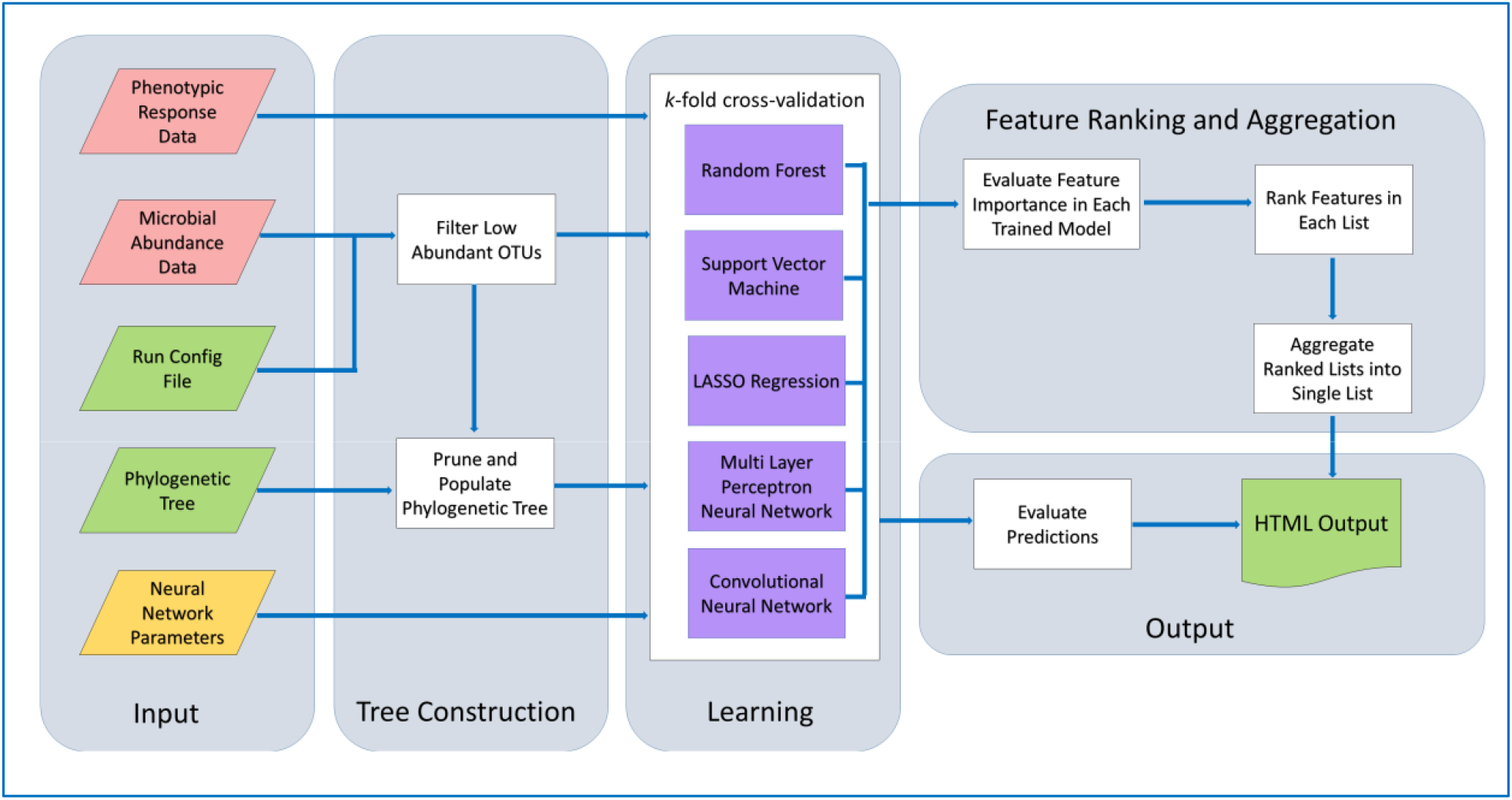
Meta-Signer Workflow. Large rounded rectangles represent different modules of the work ow. Red inputs are required and provided by the user. Green inputs are required but included in the package. Yellow inputs are optional and will be generated and saved if not provided

### Input

Meta-Signer uses relative taxa abundance data and phylogenetic taxonomy information to construct a taxonomic tree. The input to Meta-Signer are (1) a tab separated file of taxa relative abundance values where each row represents an taxon and each column represents a sample, and (2) a line separated list of response values where each row represent the phenotypic response of a sample. The first column in the abundance table should be the taxonomic identification of the taxon. Each level of taxonomy should be represented using the first letter of the taxonomic level (k, p, c, o, f, g, s), followed by two underscore characters, and then the taxonomic identification at that level. Two additional required files are (3) the run configuration file with user specified parameters, and (4) a phylogenetic tree of all microbes in newick format. Templates for both of these files are provided. A final optional file that Meta-Signer will use is (5) model parameters for the neural network architectures in JSON format. If this file is not found, Meta-Signer will tune the parameters and save them for later use.

### Tree Construction Module

The taxa relative abundance data are first pre-processed and normalized. To do so, abundance values are converted into relative abundance by dividing each value by the sum of abundance values in its respective sample. Next, low abundant taxa are filtered out based on a proportion threshold specified by the user in the configuration file. A taxon is removed if it is not present at the specified proportion level either overall or within a single class. The remaining abundance values are then log-transformed and normalized based on a user specified method (Z-normalization or min-max normalization).

Next, a taxonomic tree specific to the dataset is generated by pruning a tree template using the remaining taxon across all samples in the dataset. A tree template generated using PhyloT [28] is provided in the package. The nodes of the pruned tree are then populated with abundance values such that an internal node’s abundance is the sum of the abundance values of its children. An example of the populated tree is shown in Fig. 2 and the algorithms are shown in Fig. S1 and S2 in Additional File 1.

**Figure 2.**
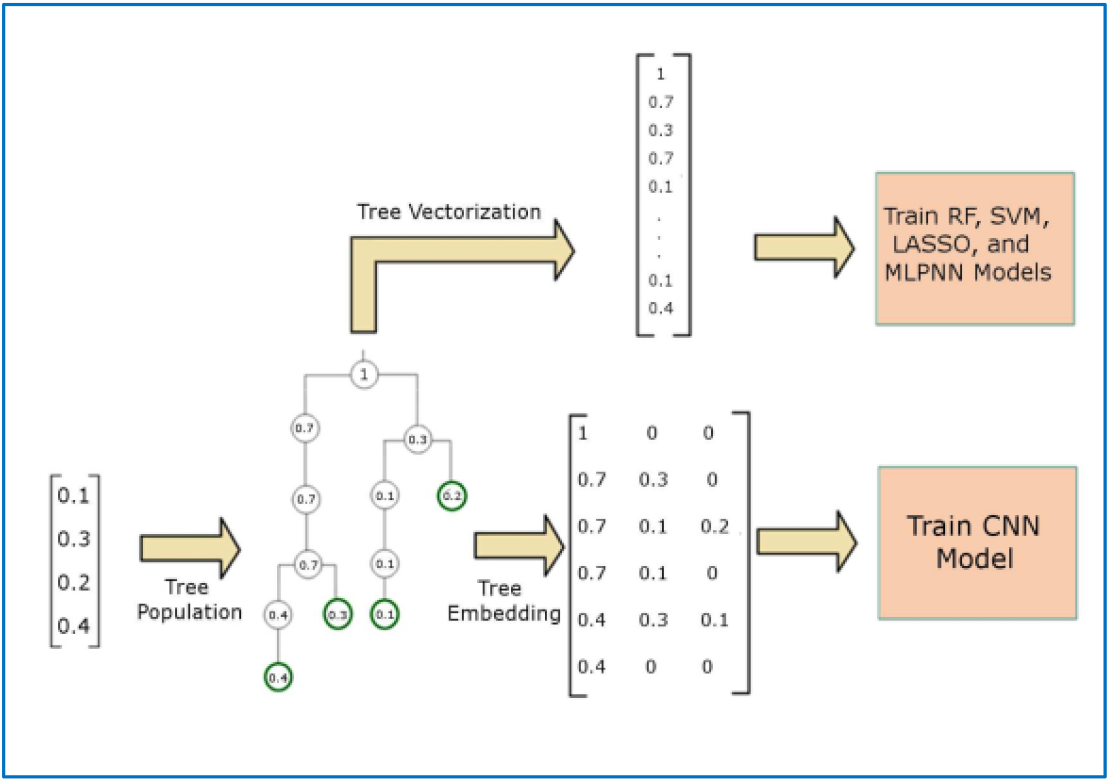
Generating tree-level taxa. A vector of relative taxa abundance is used to populate a taxonomic tree. This tree can either be represented as a matrix to be used as an input to PopPhy-CNN or be vectorized and used with any other ML method.

### Learning Module

Meta-Signer includes three classic ML models (RF, Linear SVM, LASSO), and two DNN models (MLPNN, PopPhy-CNN) in its Learning Module. RF models are decision tree learning models that are trained in an ensemble fashion, taking the average of the ensemble to give a robust decision tree [29].

While growing each tree, a decision is made at each node by selecting the feature from a random subset of features that best splits the data into two subsets based on Gini impurity of each subset. Given a set of items with *k* classes, let *p_i_* be the proportion of samples of class *i* for *i* ∈ {1, …,*l*}. The Gini impurity of the set is calculated as

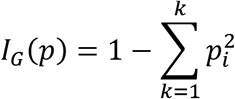

SVMs are supervised machine learning models that try to learn the best hyperplane that separates two classes of data [30]. In case of linear SVMs, from the hyperplane we can obtain a set of weights, *w*, and an intercept, *b*. The class of the sample *x_i_*, can then be determined as

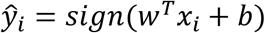

LASSO regression is a form of least squares regression that uses shrinkage to reduce the total number of model parameters in the final model [31]. This is achieved using L1 regularization in order to penalize the absolute value of the weights, eliminating a portion of the weights to create a sparse model. Given a set of samples *X* = {*x*_1_, …*x_n_*}, where each sample has *m* features and response values *Y* = {*y*_1_, …, *y_n_*}, the model minimizes the cost

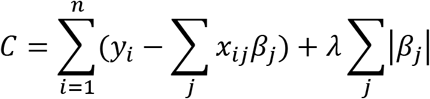

where *β_j_* are the weight parameters which are penalized with the regularization parameter *λ*.

Neural networks are consisted of multiple layers of nodes that are fully connected with edges constituting weights [32]. The values of a hidden layer are a linear combination of the values from the previous layer which is passed through a non-linear activation function. More explicitly, the values of a hidden layer *h_ι_* is calculated as

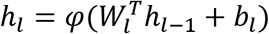

where *h*_*ι*–1_ are the values from the previous hidden layer, *W_ι_* are the weights connecting *h*_*ι*–1_ to *h_ι_*, *b_ι_* is a bias value, and is a non-linear activation function.

PopPhy-CNN was originally designed by our group [25, 26]. PopPhy-CNN explores relationship between taxa by treating a populated taxonomic tree as a type of image. Our model is based on a CNN architecture which utilizes shared kernel weights to learn relationships in the data by constructing multiple feature maps. More explicitly, given an input matrix *M*, a kernel *k* with a set of weights *w*^(*k*)^ ∈ *R^r x s^*, the velocity at position (*i,j*) of the feature map is defined as:

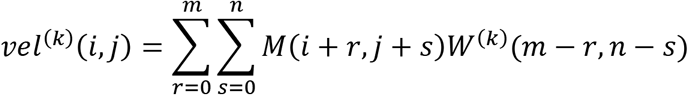

The velocities of a feature map are passed through a non-linear activation and then max pooling is applied to the feature map. More detail about PopPhy-CNN and the other learning models is given in Additional File 1.

Before training, Meta-Signer checks for a file containing network hyper-parameters for MLPNN and CNN models. If it does not find this file, Meta-Signer will use the first partition of the cross-validation to empirically determine the hyper-parameters. This is done using another cross-validation on the training set of the first partition. In addition, using the configuration file, the user can set custom parameters and even disable any of the learning models if they do not wish to incorporate it into their results. In Meta-Signer, RF, SVM, and LASSO are trained using the *scikit-learn* python package and MLPNN and CNN are trained using *Tensorflow*.

### Feature Ranking Module

For each ML model in each cross-validated partition, Meta-Signer extracts the feature scores and uses the scores to construct a ranked feature list. RF features were scored using a method called mean decrease impurity. For each node, the importance of the feature being split upon is calculated as the decrease in Gini impurity from before and after the split. This value is then weighted by the proportion of total samples that were split upon that node. A feature’s importance is then calculated by averaging the weighted importance values of nodes that split using that feature across all trees in the ensemble. Features in LASSO and SVM models were scored based on the magnitude of their weight coefficients in the decision functions.

The extraction of features from DNN models is a challenging task. We use a procedure developed in [33] to evaluate features in MLPNNs. Briefly, the MLPNN features were evaluated by calculating the cumulative weight across all layers by taking the running product of all the weight matrices in the learned networks. The method is quick and the product results in a matrix that has a column for each class and a row for each feature, and the value at a given index is that features cumulative impact for that class. We then consider a feature’s importance as the maximum impact across classes to create a single ranked list.

The novel procedure to extract features from CNNs was developed in our previous learning framework PopPhy-CNN [26]. We included the details on the full algorithm of feature importance scoring in Additional File 1 (Fig. S3 and S4) for convenience. Briefly, the procedure focused on the post analysis of the kernel map activations in the first convolutional layer prior to subsampling. It selects the top 10% of the signals from each kernel map and traces each signal back to the local area from the input from which it was generated. The local area of the input can be interpreted as a group of nodes on the tree. Each input feature’s importance is evaluated based on how much it contributed to the signal found in the kernel map.

### Feature Aggregation Module

For each partition of the cross-validation, we generate a single ranked list for each of ML model. Once the entirety of the cross-validated training is complete, the entire set of all ranked lists across all models is aggregated into a single top-*k* ranked list by minimizing the distance between the set of ranked lists and the top-*k* list, where *k* is specified by the user in the configuration file. More specifically, given a set of ranked lists {*l*_1_, …, *l_m_*}, the top-*k* ranked list, 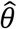, is determined as

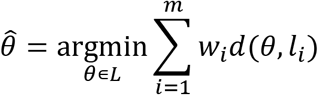

Here, *L* is the state space of top-*k* rankings, *w_i_* is a weight associated with *l_i_*, and *d*(*θ, l_i_*) is the distance between a proposed top-*k* ranked list, *θ*, and *_i_*. We consider original and tree-level taxa separately, resulting in two separate top-*k* lists.

The aggregation is performed using the R package *RankAggreg* [27]. This package uses a cross-entropy based approach with Markov Chain Monte Carlo sampling to find the top-*k* features that minimize the sum of the distances between each of the input sets and the generated top-*k* set. The distance used is the Spearman’s Correlation. Each input ranked list is weighted in the aggregation by the area under the receiver operating curve (AUC) for binary classification or the generalized Matthews correlation coefficient (MCC) for multiclass classification. When using MCC, negative values are clipped to 0 in order to prevent negative weights.

### Output

After the model predictions are evaluated and the features are ranked into a single list, Meta-Signer provides a summary of the results in a portable HTML file. The file contains a description of the run and evaluation metrics for the different models in the form of both a table and boxplots. It also provides the distribution of the feature (taxa) scores for each machine learning model. Lastly, it provides a list of the top-*k* taxa selected from the original taxa and tree-level taxa, the proportion of individual ranking sets that each taxon was present in the top-*k*, the rank and p-value under a Wilcoxon Rank-Sum test, and the class in which the taxon was found enriched in. All images are encoded into the file, allowing the HTML file to be moved without considering the location of the images.

## Results

### Datasets used in evaluation

We used five publicly available datasets to evaluate Meta-Signer: cirrhosis, obesity, type 2 diabetes (T2D), inflammatory bowel disease (IBD), and colorectal cancer (CRC). The datasets were obtained from the MetAML package [17]. The cirrhosis dataset was taken from a study of 114 cirrhosis patients and 118 healthy subjects [34]. The T2D dataset was a combination of two studies [5, 35] yielding a total of 223 patients with T2D and 217 healthy subjects. The obesity dataset comes from a study of 292 individuals of which 89 individuals with a BMI lower than 25 kg/m^2^ were studied against 164 individuals with a BMI greater than 30 kg/m^2^ [36]. The IBD dataset included 110 subjects of which 85 were healthy, 21 had ulcerative colitis, and 4 had Crohn’s disease. Since the sample size for sub-jects with Crohn’s disease was extremely small, the two disease states were combined into a single class of 25 subjects [37]. The CRC dataset contains 134 subjects where 47 subjects are healthy, 26 subjects had small adenomas, 13 subjects had large adenomas, and 48 subjects had cancer [38]. In previous studies, subjects with small adenomas have been grouped with healthy subjects and subjects with large adenomas have been dropped. However, we wanted to evaluate our method on a multiclass dataset, so we chose to instead combine subjects with small adenomas and large adenomas into a single class of 39 subjects. The first four datasets involve binary classes and the last dataset involves three classes. A summary of the datasets used in our evaluation is shown in Table 1.

**Table 1.** Datasets. Sample sizes and numbers of taxa for datasets used for evaluation. In the case of CRC, the number of samples for each case is separated by a comma.

|  | # Case | # Control | # Taxa |
| --- | --- | --- | --- |
| Cirrhosis | 114 | 118 | 542 |
| T2D | 223 | 217 | 606 |
| Obesity | 164 | 89 | 465 |
| IBD | 25 | 85 | 443 |
| CRC | 48, 39 | 47 | 507 |

Each of these datasets was generated using Metagenomic Shotgun (MGS) sequencing. The taxa for each dataset were assigned by MetaPhlAn2, which selects taxa based on the read coverage of clade-specific markers and then estimates their relative abundance [39]. The taxa in each dataset were aggregated at the species level. For any taxon which could not be classified at the species level, we added its abundance to its genus level internal node of the tree. In our experiments, we filtered out any taxon that was not present in at least 10% of a single response class. All abundance values were then log-transformed and normalized to have a mean of 0 and unit variance.

### Evaluation of ML models

We first provide an overall evaluation of individual ML models under the same experimental protocol. This can also identify any model that has exceptionally inferior performance. If it indeed occurs then the model can be excluded from the Aggregation Module. For the multiclass CRC dataset, SVM and LASSO are evaluated using a one-verse-all approach. We use the following criteria for our model evaluation: AUC for binary classification (MCC for multiclass classification), precision, recall and F1 score. Here,

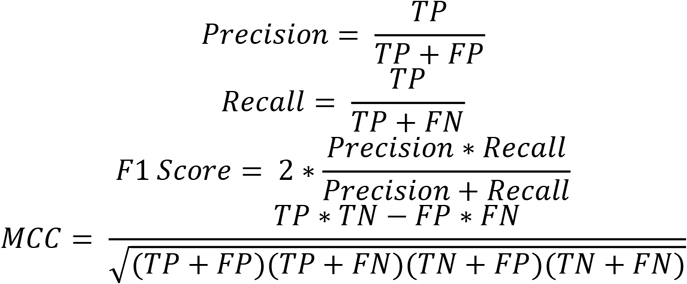

where TF, FP, TN, and FN are true positives, false positives, true negatives and false negatives, respectively.

Models were trained using 10-fold cross-validation. The average and standard deviation of each evaluating criterion from 10 times cross-validations are reported. In each 10-fold cross-validation, the dataset was partitioned into 10 sets stratified by class proportion. We then constructed 10 datasets where a single partition was left out as the test set and the remaining were used for the training set.

For RF models, a maximum of 500 trees was set and all other parameters were left as the default. Feature importance values were first obtained by training a RF using the training set and extracting the feature importance scores from the trained model. We performed 5-fold cross-validation on each training set to determine the optimal number of features to use. We evaluated models using the top 25%, 50%, 75%, and 100% of features and selected the best one. A final model was trained using the determined optimal number of features on the entire training set and evaluated on the test set.

The parameters in Linear SVM and LASSO models were determined using 5-fold cross-validation on each training set. The parameter associated with the error term in SVM was chosen from {1, 10, 100, 1000}, and the regularization parameter in LASSO model ranges from 10^-4^ to 10^-0.5^ that were space evenly on a log-scale. The best parameter value was determined using 5-fold cross-validation on the training set, and the nal model was trained on the training set and evaluated on the test set.

MLPNN and CNN models were trained for 200 epochs with a dropout rate of 0.5, a learning rate of 0.001, and an L2-regularization penalty value of 0.01. MLPNN models contained 2 layers of 32 nodes. CNN models contained a single convolutional layer with 32 square kernels of size 5 and a single fully connected layer with 32 nodes. Network hyper-parameters were found using 10-fold cross validation on the training set where 70% of the data was used to train the network, 20% of the data was used to determine when to stop early, and the last 10% was used to evaluate the model. A set of hyper-parameters was selected based on the mean performance. We only tune the network using the first cross-validation partition because we assume that the learning complexity of the cross-validation partitions is similar. We used the rectified linear unit (ReLU) activation function for hidden layers and the softmax activation function for the output layer. Each network is trained using a weighted cross-entropy loss function. For consistency in our experiments, we shared the tuned parameters of one dataset, in our case the cirrhosis dataset, between datasets, believing that the complexity of the classification tasks was comparable between datasets.

In our experiments, we observed that RF models performed the best overall compared to the other ML models (Table 2) using the tree taxa as features. The performance using the original taxa had a similar pattern (Table S1 in Additional File 1). The MLPNN and PopPhy-CNN models showed improved performance over SVM and LASSO, but overall they were not superior to RF (Table 2). However, we observed that within a cross-validated evaluation there were partitions in which RF models were outperformed by other models (Fig. 3). Similar comparisons between all models are shown in Fig. S5 in Additional File 1. Therefore, we decided to use features from multiple learning models across the multiple partitions. In addition, we noticed a slight decrease in performance when using the entire set of tree-level features. Despite the reduction in performance, we believe that including the tree taxa can help provide insight to features associated with dysbiosis at multiple levels of the taxonomic tree

**Figure 3.**
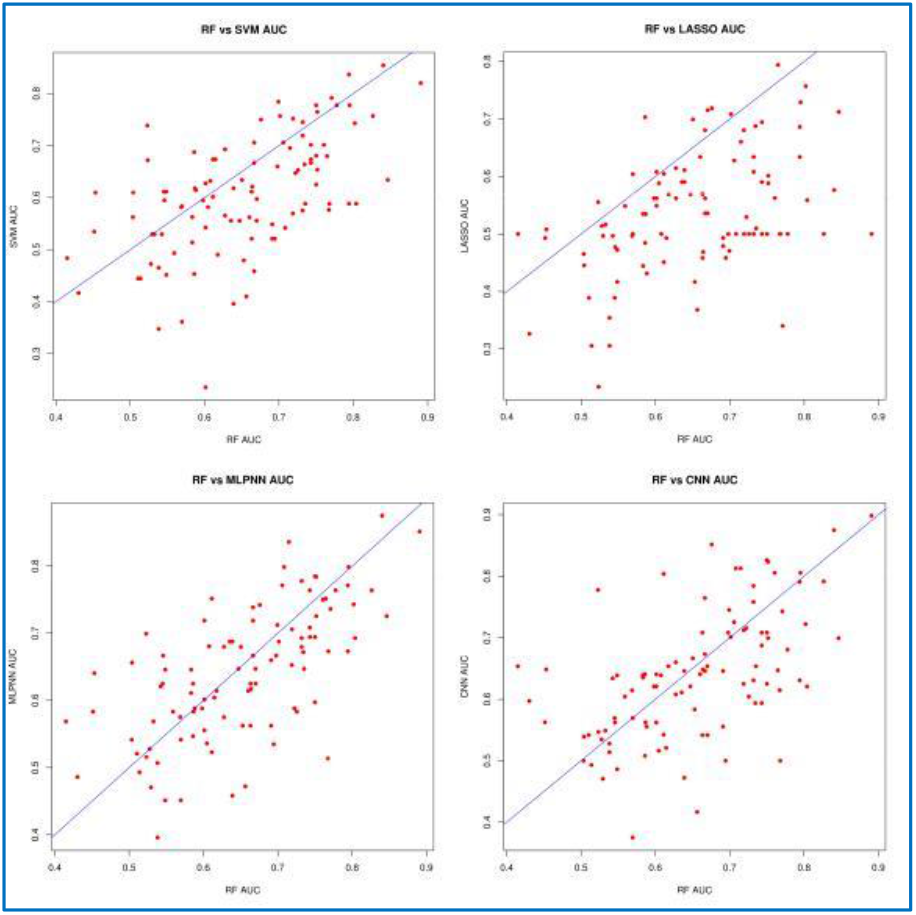
Scatterplots AUC values of RF compared to those of SVM, LASSO, MLPNN, and CNN models. Each plot represents the results of 10 iterations of 10 fold cross-validation using the tree taxa of the obesity dataset. Each point represents a single partition. Points above the blue line are instances in which the other method outperforms RF.

**Table 2.** Performance of the individual ML models using tree taxa. Mean AUC, MCC, Precision, Recall, and F1 score are shown for all five datasets using the different machine learning methods. Standard deviation is shown in parentheses

|  |  | RF | SVM | LASSO | MLPNN | PopPhy-CNN |
| --- | --- | --- | --- | --- | --- | --- |
| Cirrhosis | AUC | **0.937 (0.057)** | 0.827 (0.083) | 0.916 (0.065) | 0.914 (0.069) | 0.924 (0.063) |
|  | MCC | **0.767 (0.123)** | 0.516 (0.186) | 0.622 (0.159) | 0.680 (0.135) | 0.685 (0.153) |
|  | Precision | **0.890 (0.059)** | 0.764 (0.094) | 0.820 (0.081) | 0.850 (0.067) | 0.852 (0.075) |
|  | Recall | **0.878 (0.065)** | 0.753 (0.093) | 0.802 (0.080) | 0.830 (0.069) | 0.833 (0.080) |
|  | F1 | **0.876 (0.066)** | 0.751 (0.094) | 0.799 (0.081) | 0.828 (0.071) | 0.830 (0.082) |
| T2D | AUC | **0.738 (0.058)** | 0.610 (0.078) | 0.676 (0.074) | 0.695 (0.064) | 0.702 (0.067) |
|  | MCC | **0.326 (0.094)** | 0.153 (0.135) | 0.239 (0.129) | 0.283 (0.117) | 0.277 (0.125) |
|  | Precision | **0.665 (0.047)** | 0.578 (0.069) | 0.621 (0.065) | 0.644 (0.059) | 0.641 (0.064) |
|  | Recall | **0.662 (0.046)** | 0.575 (0.066) | 0.619 (0.064) | 0.639 (0.057) | 0.637 (0.061) |
|  | F1 | **0.660 (0.047)** | 0.572 (0.066) | 0.617 (0.065) | 0.636 (0.059) | 0.634 (0.062) |
| Obesity | AUC | **0.648 (0.116)** | 0.580 (0.120) | 0.563 (0.096) | 0.629 (0.105) | 0.634 (0.105) |
|  | MCC | 0.038 (0.182) | 0.018 (0.158) | 0.029 (0.187) | 0.185 (0.179) | **0.186 (0.162)** |
|  | Precision | 0.555 (0.123) | 0.510 (0.137) | 0.534 (0.140) | 0.630 (0.084) | **0.632 (0.078)** |
|  | Recall | **0.631 (0.051)** | **0.635 (0.049)** | 0.631 (0.059) | 0.624 (0.080) | 0.618 (0.076) |
|  | F1 | 0.556 (0.068) | 0.532 (0.054) | 0.545 (0.069) | **0.622 (0.080)** | 0.616 (0.075) |
| IBD | AUC | **0.890 (0.103)** | 0.801 (0.166) | 0.812 (0.149) | 0.839 (0.145) | 0.856 (0.125) |
|  | MCC | 0.339 (0.340) | 0.179 (0.324) | 0.224 (0.318) | 0.432 (0.277) | **0.503 (0.290)** |
|  | Precision | 0.768 (0.146) | 0.706 (0.136) | 0.715 (0.139) | 0.810 (0.107) | **0.831 (0.114)** |
|  | Recall | **0.823 (0.073)** | 0.765 (0.093) | 0.795 (0.073) | 0.778 (0.106) | **0.823 (0.097)** |
|  | F1 | 0.775 (0.101) | 0.723 (0.102) | 0.741 (0.097) | 0.779 (0.101) | **0.816 (0.101)** |
| CRC | AUC | - | - | - | - | - |
|  | MCC | **0.338 (0.215)** | 0.318 (0.184) | 0.223 (0.187) | 0.239 (0.186) | 0.240 (0.226) |
|  | Precision | **0.552 (0.144)** | 0.536 (0.157) | 0.439 (0.196) | 0.511 (0.140) | 0.515 (0.148) |
|  | Recall | **0.554 (0.138)** | 0.539 (0.115) | 0.461 (0.100) | 0.488 (0.118) | 0.488 (0.145) |
|  | F1 | **0.533 (0.132)** | 0.510 (0.123) | 0.381 (0.126) | 0.476 (0.121) | 0.477 (0.138) |

### Taxa extracted from Meta-Signer are more informative than other methods

To evaluate Meta-Signer, we benchmark against other three methods: two previously published methods: Biosigner [40], a hierarchical feature engineering (HFE) method [41], and a non-parametric Wilcoxon rank-sum test. Biosigner is a generic ML driven feature selection method for omics data and available in R. It uses trained RF, SVM and Partial Least Squared Discriminant Analysis models to selectively eliminate features, resulting in a single set of remaining features. HFE is implemented in Python specifically designed for metagenomic data and uses a taxonomic tree to construct hierarchical features and extracts nodes based on information gain. HFE was shown to outperform other state-of-the-art methods such as Fizzy [42] and MetaML [17]. We were not able to bench-mark against OMiAT [11], since the method requires the tree branches to contain distance values, which are not available to us.

The non-parametric Wilcoxon rank-sum test was included as a baseline method of feature ranking for comparison. We used the top 20 taxa from each method, except for Biosigner, which only identified less than 20 taxa in every dataset evaluated.

We evaluated each method using 10-fold cross-validation. Each dataset was randomly partitioned into 10 sets, stratified to balance the class proportion. Then each method was applied to 9 of the 10 sets, the training set, to extract features. Then the taxa selected from the ML models are aggregated. The datasets were then filtered to only have the chosen taxa and models were trained on the training set and evaluated on the test set. We used SVMs with Gaussian kernels to evaluate the performance for the selected taxa from all the four methods. In order to perform fair comparison, we report the evaluation without including linear SVM models in taxa aggregation.

We observed that Meta-Signer was robust across all datasets (Table 3). In cirrhosis, T2D, and obesity, it outperformed all other models. Biosigner and HFE performed very poorly for these datasets. Biosigner often returned no taxa (in which case an AUC of 0.5 was assigned to the testing). In the cirrhosis dataset, we observed that using tree-taxa (Meta-Signer (Tree) and HFE) caused the SVM models to become less stable and predict more poorly. This could be in part that many of the tree taxa had parent-child relationships and carried redundant information. These similar taxa may have saturated the top 20 ranked taxa, leaving less room for other informative taxa. In the obesity dataset, both Biosigner and HFE extract taxa that lead to a testing AUC less than 0.5. Surprisingly, the taxa ranked from the Wilcoxon test generated the best performance for IBD. Both Wilcoxon and HFE performed well in CRC. However, although not the best in IBD and CRC, Meta-Signer was still comparable. We observe a similar pattern for performance of Meta-Signer when results of linear SVMs are included in taxa aggregation (Table S7 in Additional File 1).

**Table 3.** Comparison of Meta-Signer to other methods. Mean AUC/MCC values for the selected datasets using different feature selection methods. Standard deviation values are shown in parentheses.

|  | Meta-Signer | Meta-Signer (Tree) | Biosigner | HFE | Wilcoxon |
| --- | --- | --- | --- | --- | --- |
| Cirrhosis (AUC) | **0.905 (0.050)** | 0.699 (0.322) | 0.815 (0.231) | 0.627 (0.349) | 0.822 (0.168) |
| T2D (AUC) | **0.634 (0.069)** | 0.629 (0.080) | 0.578 (0.052) | 0.584 (0.098) | 0.563 (0.138) |
| Obesity (AUC) | **0.600 (0.135)** | 0.529 (0.131) | 0.437 (0.103) | 0.485 (0.127) | 0.582 (0.189) |
| IBD (AUC) | 0.794 (0.159) | 0.792 (0.143) | 0.639 (0.111) | 0.831 (0.091) | **0.875 (0.100)** |
| CRC (MCC) | 0.325 (0.135) | 0.284 (0.185) | - | **0.352 (0.161)** | 0.246 (0.208) |

We then looked at the overlap of original taxa extracted from Meta-Signer to those extracted using a Wilcoxon test and Biosigner. HFE and Meta-Signer using the tree taxa were excluded since they both created their own unique feature space and were not comparable. To compare methods, each method was applied to the entire dataset, generating a single ranked list for each method. We then computed the overlap for the top 20 taxa from Meta-Signer, the Wilcoxon test, and Biosigner. Since Biosigner does not run on multi-class datasets, is not considered in the overlap of the CRC dataset. The results are shown Fig. 4 and the top 20 taxa identified in each dataset by Meta-Signer are shown in Tables S2–S6 in Additional File 1. In general, Meta-Signer and the Wilcoxon test have the largest overlap for identified taxa.

**Figure 4.**
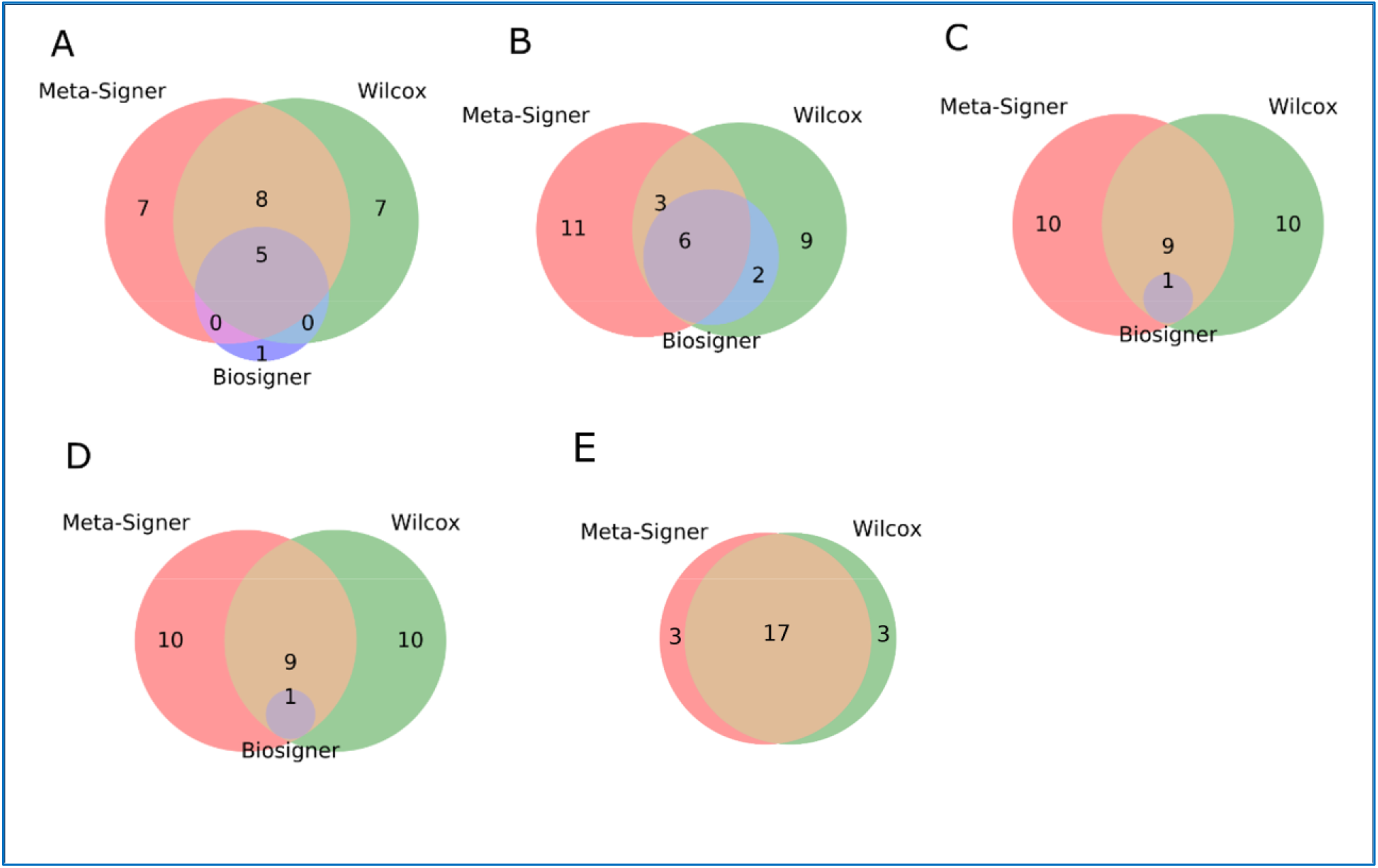
Taxa overlap. Taxa overlap between Meta-Signer, Wilcoxon test, and Biosigner. Taxa overlaps are shown for (A) cirrhosis, (B) T2D, (C) obesity, (D) IBD, and (E) CRC datasets.

Finally, we examined at the proportion of how often the extracted taxa appeared in the top 20 taxa in each model independently in Meta-Signer. We observed a large consensus in many taxa, however, a few taxa were missed by some models completely and picked up by the others (Tables S8 and S9 in Additional File 1). In addition, we observed that PopPhy-CNN was not contributing as much as the other methods to the top 20 selected taxa in each dataset. Upon visual inspection, we observed that the taxa from the CNN models were more often at higher taxonomic levels (class and family levels and super levels). Therefore, we suspected that it captured information at higher aggregated taxa, and as such, less number of taxa are required. To investigate this, we trained SVM models using just the top 10 taxa after aggregating the ranked lists from the CNN models only and compared the performance to Meta-Signer on the tree taxa. Except for the IBD dataset, the taxa extracted just from the PopPhy-CNN models are comparable or even superior to the ensemble aggregated taxa. This makes us believe that PopPhy-CNN is finding useful taxa, however, they are being dampened by the other models in the ensemble aggregation. A table comparing the top 10 taxa of the PopPhy-CNN model and Meta-Signer is shown in Table S10 in Additional File 1.

### Taxa extracted from Meta-Signer are biologically relevant - A case study

We focus on the obesity data to examine the relevance of the extracted taxa from Meta-Signer (Fig. 5). A number of them have been identified as relevant to obesity from previous studies. We observed an enrichment of *Ruminococcus torques* and *Bacteroides ovatus* in obese subjects, both of which have been shown to be positively correlated with metabolic syndrome traits [43, 44]. *Clostridium leptum* was also found to be enriched in obese subjects, which agrees with the findings of a previous study in Danish infants [45], however there have been conflicting observations on the correlation of this species with regards to obesity [46]. Meta-Signer was also able to identify *Dorea formicigenerans* and *Mitsuokella* as enriched in obese patients which has been reported in previous studies [44, 47] as well as *Megasphaera elsdenii* which has been identified at a genus level to be associated with obesity [48]. Additionally, we observed that *Oscillibacter* was enriched in obese subjects, which has been associated to the consumption of high-fat diets [49]. Interestingly, *Ruminococcus torques*, *Bacteroides ovatus*, *Mitsuokella*, and *Megasphaera elsdenii* are insignificant based on the Wilcoxon rank test. Two species enriched in lean patients were *Ruminococcus lactaris*, a species found enriched in human patients after being treated for metabolic syndrome using resistant starch type 4 [50], and *Ruminococcus avefaciens*, a species found to be enriched in non-obese mice [51]. *Ruminococcus lactaris* is also insignificant from the Wilcoxon rank test.

**Figure 5.**
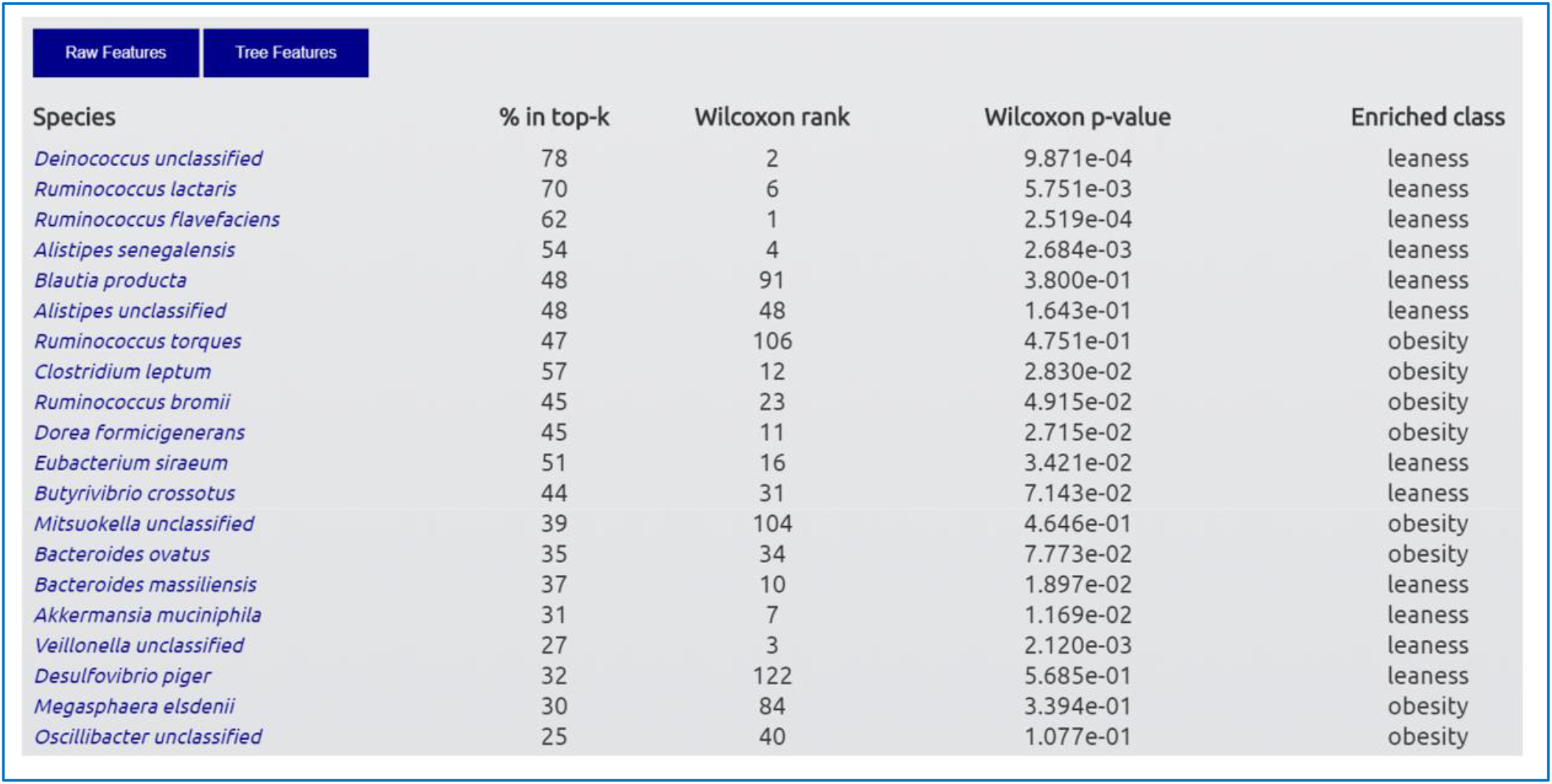
Top 20 taxa selected in the obesity dataset. The output HTML file provides the feature list, the percent of individual ranked lists across all trained models in which the feature was found in the top-k, the feature rank and p-value based on the Wilcoxon test, and the class that the feature is enriched in.

In addition, we found that microbes from the genera *Alistipes* as well as *Akkermansia muciniphila* were enriched in lean patients. Microbes from these genera have been associated to successful weight loss in obese patients after a regiment of a low-calorie diet, exercise, and behavioral therapy [52, 53]. We also identified *Ruminococcus bromii*, *Eubacterium siraeum*, and *Butyrivibrio crossotus* to be enriched in lean subjects, which have all been observed in previous studies [54, 55, 56]. Another taxon that was captured strongly and at multiple levels in the tree was *Deinoccocus*. This genus has not been strongly implicated with obesity, however, a recent study has shown that some species of this genera create amylosucrase. The study showed that modifying consumed chestnut starch with this enzyme lead to a suppression of insulin signaling and a reduction of fat accumulation [57].

Lastly, we wanted to visualize the landscape identified by Meta-Signer. To do this, we combined the ranked original taxa and ranked tree taxa into a single set and isolated the subtree containing this set of taxa. We annotated the subtree to visualize which branches were enriched in both lean and obese individuals. We connected nodes using a solid line of the same color if Meta-Signer identified nodes at higher taxonomic levels along the branch. Otherwise, we connected nodes using a dashed line of the same color until a branch occurred. The visualized subtree is shown in Fig. 6.

**Figure 6.**
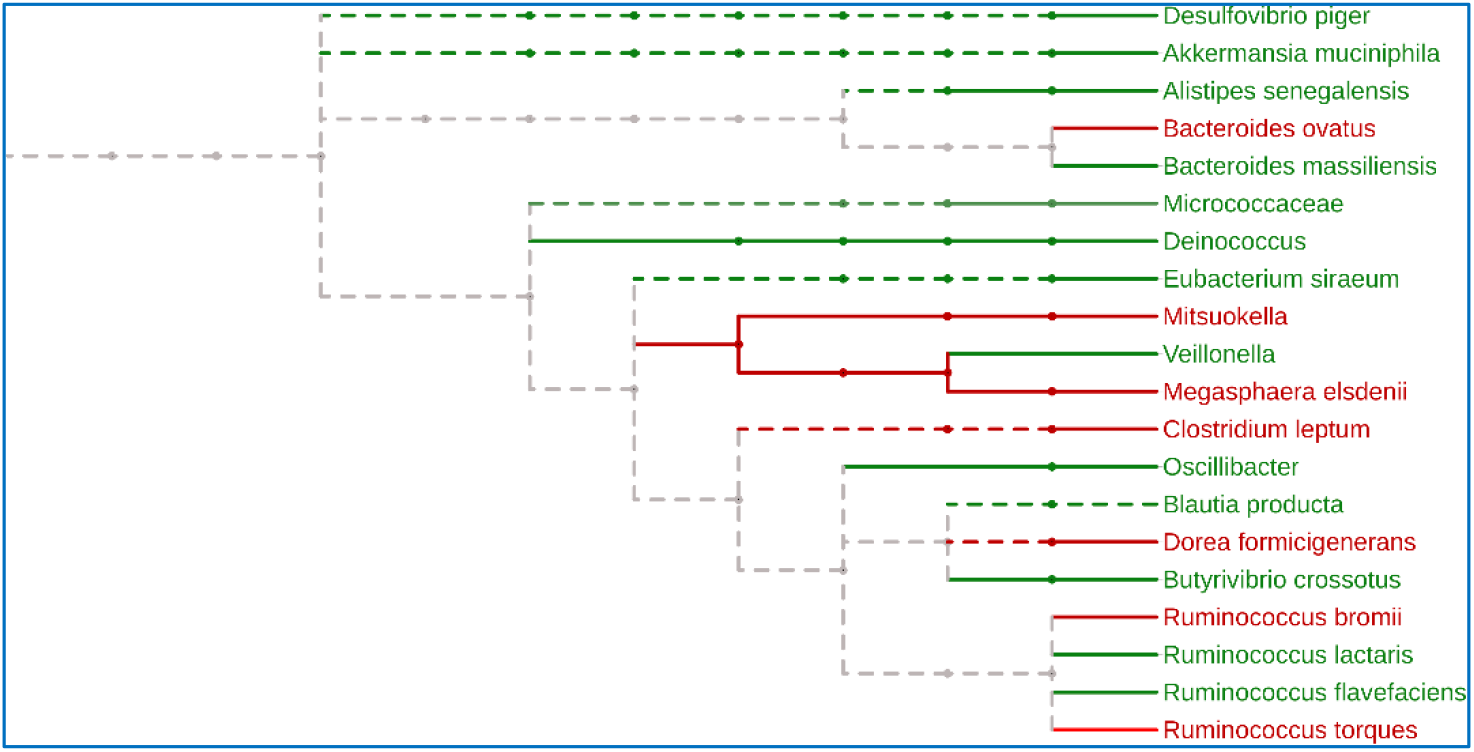
Subtree of identified taxa in the obesity dataset. A taxonomic tree of the combined ranked tree and original taxa for lean (green) and obese (red) subjects. Solid lines indicate taxonomic levels identified using the tree taxa. Dashed lines are used to extend phenotypic associations through the tree at nodes that were not identified until a branch occurs.

## Discussion

We have developed Meta-Signer, a user-friendly tool for the extraction of robust microbial taxa that are predictive to host phenotype at multiple taxonomic levels from multiple machine learning models. Meta-Signer is able to leverage biological knowledge in microbial taxa relative abundance profiles through a taxonomic tree by our novel propagation and matrix-representation procedure. By training different types of machine learning models, Meta-Signer exploits the similarities in the ranked lists of taxa learned by individual machines learning models to create a single aggregated set of informative microbial taxa for host phenotype prediction.

Using five metagenomic datasets, we demonstrated that Meta-Signer not only outperforms state-of-the-art methods of feature ranking but also is robust. In addition, in our case study on the obesity dataset, we have shown that the many of the taxa extracted from Meta-Signer have been found biologically relevant in previous studies. It should be emphasized that the objective of the current work is not aiming at the establishment of the highest predictive model, but at the identification of the most robust set of taxa that are predictive to the host phenotype. The identified taxa could be used for follow-up validation [17].

There are several limitations to our method. The first is that there is no predetermined way to know how many taxa to extract. We set a default of 20 taxa, but in some cases this may be too many and in others it may not be enough. This is compounded with the fact that the tree-level taxa often contain redundant information since both child and parent nodes can be very similar or even equal in abundance. This may cause both features to be selected, leading to the feature list to become saturated with that branch of the tree, reducing the amount of space for features which can bring in new information. Lastly, deep learning models require a GPU to be time efficient. However, we do allow users to turn which models they want to train o and on in case they do not want to run them.

There are a few directions for further improvements. First, we plan to incorporate ways to reduce the saturation caused when using tree-level taxa. This would allow a more diverse set of features when using the taxonomic tree. Second, we currently drop any features not found when constructing and pruning the tree. This could lead to a loss of information, and these features should still be integrated into the tree in a meaningful way

## Conclusion

In conclusion, Meta-Signer is a user-friendly tool to identify a robust set of highly informative microbial taxa that are predictive of human disease status from a metagenomic dataset.

## Abbreviations

ML: machine learning;
RF: random forest;
SVM: support vector machine;
LASSO: least absolute shrinkage and selection operator;
DNN: deep neural network;
MLPNN: multi-layer perceptron neural network;
CNN: convolutional neural network;
AUC: area under curve;
MCC: Matthews correlation coefficient;
T2D: type 2 diabetes;
HFE: hierarchical feature engineering

## Declarations

### Ethics approval and consent to participate

Not applicable

### Consent for publication

Not applicable

### Availability of data and material

The software code and data are publicly available at https://github.com/YDaiLab/Meta-Signer.

### Author’s contributions

DR and YD conceived the idea, developed the methods and designed the evaluation plan. DR implemented the package and conducted evaluation. DR, YD, AM, and JS participated the result analysis and wrote the manuscript. All authors read and approved the final manuscript.

## Acknowledgements

We gratefully acknowledge the support of NVIDIA Corporation with the donation of the Titan Xp GPU used for this research.

## Additional file

### Tree Construction

In our work, the online tool PhyloT was used to create the phylogenetic tree, and a constant distance of one between nodes in the tree is assumed. This procedure was developed in our previous work PopPhy-CNN. For the sake of completeness, we included here.

This tree is essentially a taxonomic tree. The phylogenetic tree is structured using ancestral nodes from both taxonomic groups and subgroups with no defined distances between nodes. Therefore, we define the distance between any two nodes by the number of nodes between them. The taxonomic tree is used as a template to construct a populated tree for each sample in the dataset. In order to populate a tree, the abundance value of each taxon from a sample is assigned to its respective node in the tree. The tree is then populated such that an abundance value for each internal node is equal to the sum of its children’s abundance values. The algorithm is shown in Fig. S1.

**Figure S1.**
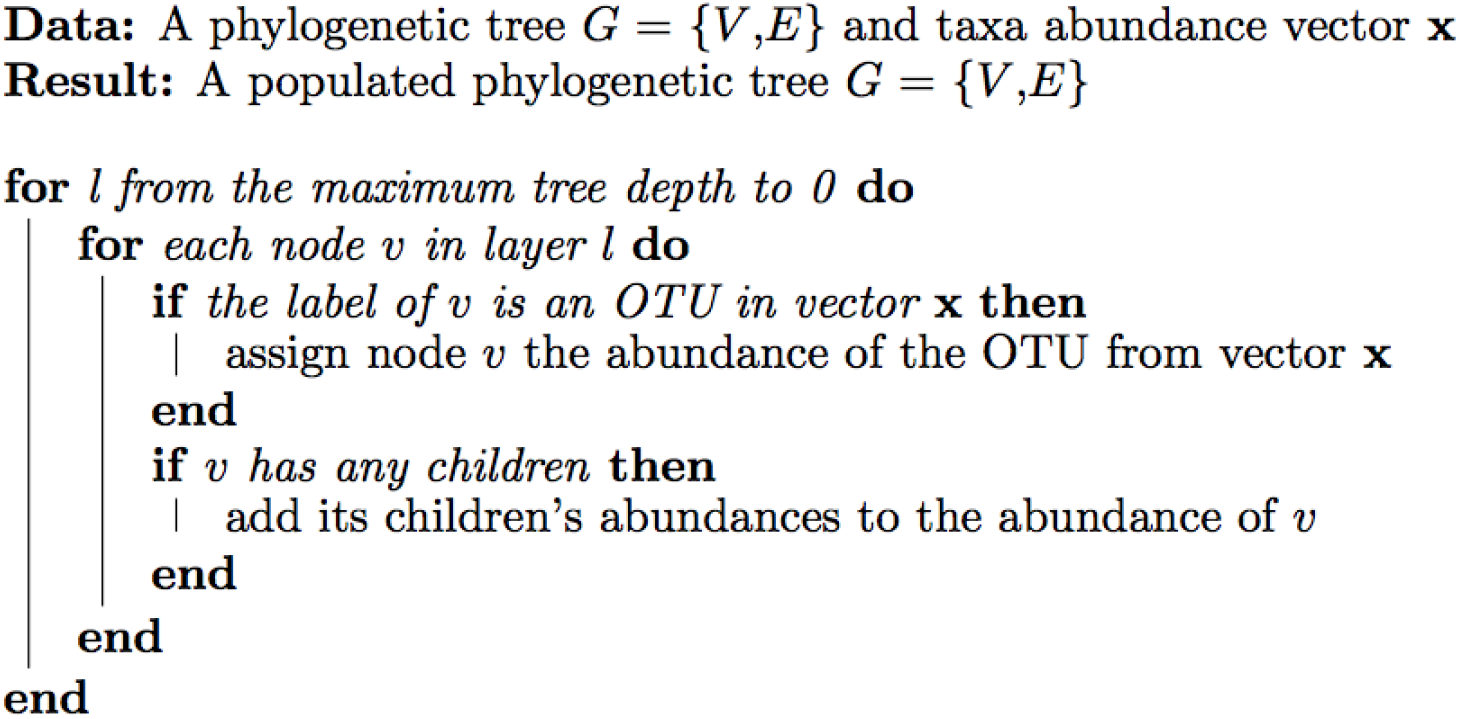
Algorithm for tree population

Since Convolutional neural networks (CNNs) are very successful in image processing where inputs are a multi-dimensional matrix, we transform the populated tree into a matrix format that contains meaningful similarity information both vertically through the rows and horizontally through the columns. We began by combining the taxa abundances and the taxonomy tree by assigning the nodes corresponding to the taxa their respective abundances. This is followed by populating the rest of the tree where a parent node’s abundance is the sum of its children’s abundances. This is preformed from the bottom upwards to the root node, which is populated with the sum of the abundance from all organisms found in the community. The algorithm for tree-matrix representation is outlined in Fig. S2.

**Figure S2.**
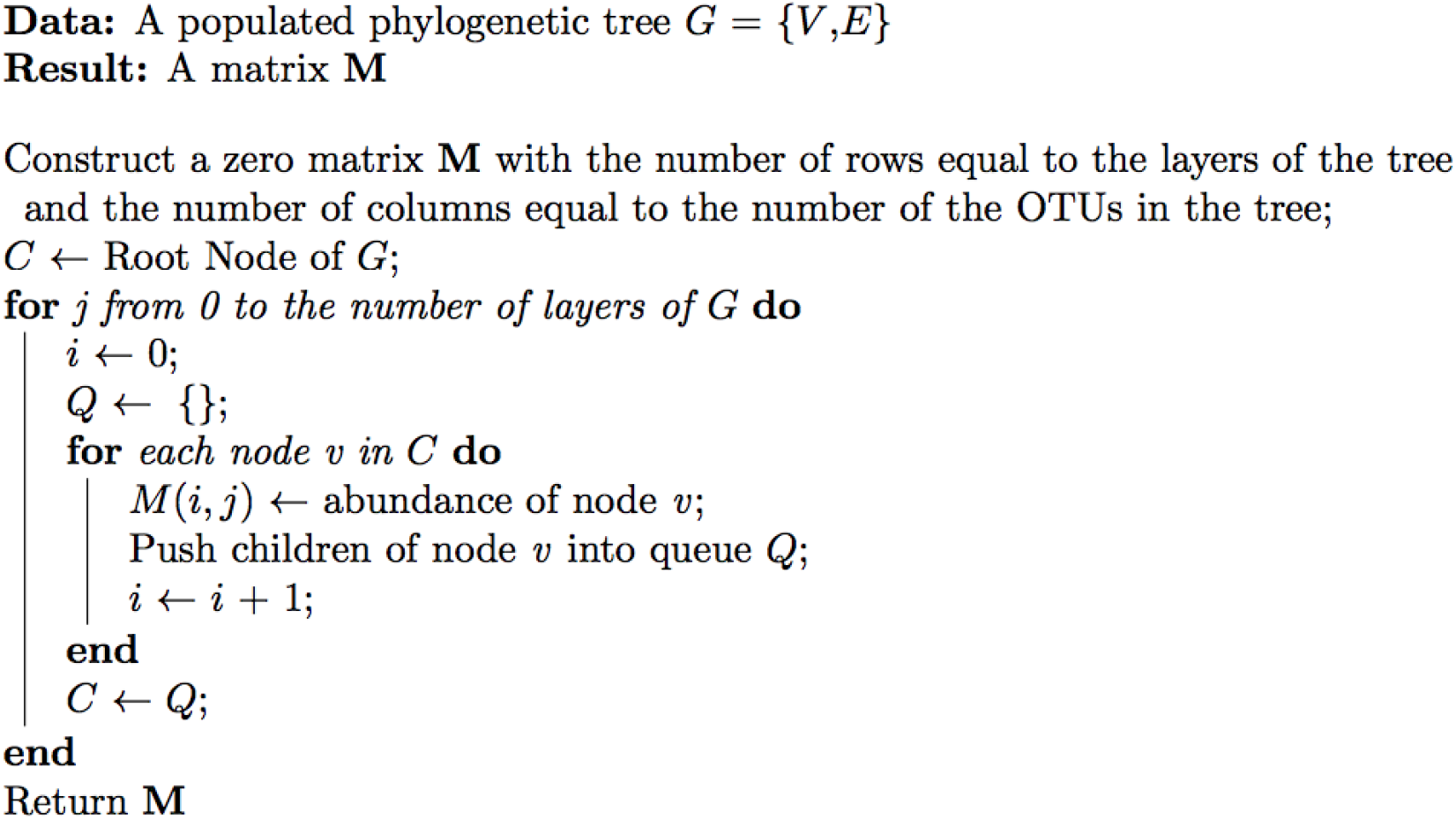
Algorithm for tree-matrix representation

These matrices are then used to train CNN models. In order to use these hierarchical features to train other machine learning models, we vectorized the tree nodes. To do so, we build up a vector, beginning with the root of the tree. Then, for each hierarchical layer of the tree, moving from the top to the bottom, the nodes from that layer are sorted by name and then appended to the vector, resulting in a of length |**V**|, where **V** is the set of nodes in the tree.

### Machine Learning Models

#### Random Forest

Random forest (RF) models are decision tree learning models that are trained in a supervised manner. The model trains an ensemble of trees and takes the average of the ensemble to give a robust decision tree. While growing each tree, a decision is made at each node by selecting the best feature from a random subset of features that best splits the data into two subsets. In our method, we generate decision rules using the Gini impurity metric. Given a set of items with *k* classes, let *p_i_* be the proportion of samples of class *i* for *i* ∈ {1 … *k*}. The Gini impurity of the set is calculated as

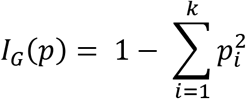

Our method implements the RF model using the *scikit-learn* python library. Once trained, features are then extracted by evaluating the mean decrease impurity. For each node, the importance of the feature being split upon is calculated as the decrease in Gini impurity from before and after the split. This value is then weighted by the proportion of total samples that were split upon that node. A feature’s importance is then calculated by averaging the weighted importance values of nodes that split using that feature across all trees in the ensemble.

#### Support Vector Machines

Support Vector Machines (SVMs) are supervised machine-learning models that try to learn the best hyperplane that separates two classes of data. SVMs, in particularly, linear SVMs, try to find the hyperplane that maximizes the margin, or separation distance, between the two classes. The orientation and position of the hyperplane is driven by a subset of data points called support vectors that lie close to the hyperplane. In case of linear SVM, from the hyperplane we can obtain a set of weights, *w*, and an intercept, *b*. The class of the sample *x_i_* can then be determined as

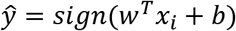

Since the above function can be used to rank the importance of features, we used the linear SVMs in Meta-Signer for feature extraction. To evaluate Meta-Signer and other exist methods, we used the nonlinear SVMs with the radial basis function kernel, 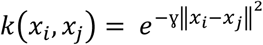 in order to achieve good performance. In our method, we allow a grid search over two kernels using the *scikit-learn* python library.

#### LASSO Regression

Least Absolute Shrinkage and Selection Operator (LASSO) regression is a form of least squares regression that uses shrinkage to reduce the total number of model parameters in the final model. This is achieved using L1 regularization in order to penalize the absolute value of the weights, eliminating a portion of the weights to create a sparse model. Given a set of samples *X* = {*x*_1_;*x*_2_, …, *x_n_*} where each sample has *m* features and classes *Y* = {*y*_1_, *y*_2_,… *y_n_*}, the model minimizes the cos

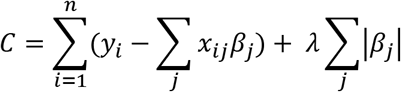

where *β* are the weight parameters which are penalized with the regularization parameter *λ*. As *λ* increases, more of the model parameters are shrunk to zero and the model becomes sparser. In order to extract the features from LASSO models, we consider positive weights to be applied to features significant to the positive class and negative features to be significant to the negative class. We rank features from a LASSO model based on the magnitude of their corresponding *β* value.

#### Multi-Layer Perceptron Neural Network

Neural networks are consisted of multiple layers of nodes that are fully connected with edges constituting weights. The values of a hidden layer are a linear combination of the values from the previous layer which is passed through a non-linear activation function. More explicitly, the values of a hidden layer *h_ι_* is calculated as

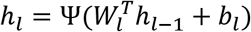

where *h*_*ι*-1_ are the values from the previous hidden layer, *W_ι_* are the weights connecting *h*_*ι*-1_ to *h_ι_*, *b_ι_* is a bias value, and Ψ is a non-linear activation function. This non-linear transformation is applied over multiple layers. In our method, we use the Rectified Linear Unit (ReLU) activation function. The ReLU activation function sets all negative values equal to 0 and all positive values are unchanged. The output layer uses the softmax activation function to predict class probabilities.

Once a model is trained, we evaluate the importance of each feature by calculating it’s cumulative weight contribution for each class. The larger the value of the cumulative contribution, the larger the influence a change in a feature will have on the prediction. More explicitly, the importance matrix can be calculated as

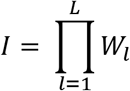

which will result in matrix *I* ∈ ℝ^*p×c*^ where *p* is the number of features and *c* is the number of classes. Features were then ranked by the maximum value across all the classes to give a single ranked list.

#### Convolutional Neural Network (PopPhy-CNN)

Convolutional neural networks (CNNs) are a deep learning model that consider groups of local features in machinelearning tasks. They have become popular in image processing as well as natural language processing. We have developed a novel CNN framework based on the input to exploit taxonomic information of the taxa by treating a populated taxonomic tree as a type of image. Our framework (PopPhy-CNN) contains one or more convolutional layers, in which a set of kernels are used to capture various signals and generate a set of feature maps. This is usually followed with at least one fully connected hidden layer, followed lastly by the output layer. A generic structure of a CNN is shown in Fig. S3.

**Figure S3.**
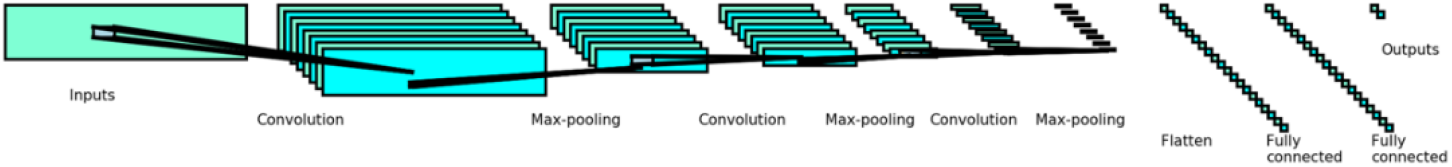
A generic CNN structure.

Given an input matrix *M*, a kernel *k* with a set of weights *W*^(*k*)^ ∈ ℝ^*r×s*^, the velocity at position (*i,j*) is defined as:

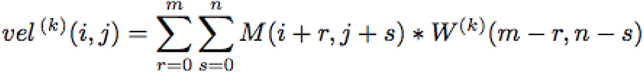

This velocity value is then passed to a non-linear activation function and pooling is performed over the entire set. In PopPhy-CNN, we use max pooling and the ReLU activation function for convolutional and fully connected layers. In the output layer, we use a softmax activation in order to predict the class probabilities, assigning the predicted class to the highest probability. We train the entire model using stochastic gradient descent (SGD) with the cost of a sample with class *c* defined as:

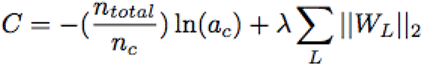

Where *n_total_* is the total number of samples, *n_c_* is the number of samples with class *c, a_c_* is the predicted probability using the softmax activation of class *c*, and *λ* is a regularization penalty coefficient for an L2-normalization of the weights.

#### Feature scoring in PopPhy-CNN

To evaluate features from a trained CNN model, we propose here a scoring schedule to extract learned features. We focus on the post analysis of the map activations in the first convolutional layer prior to subsampling. This allows to evaluate which positions in the input contribute most to the highest activations in the learned CNNs. A visualization of the generation for a feature map for the first layer is shown in Fig. S4.

**Figure S4.**
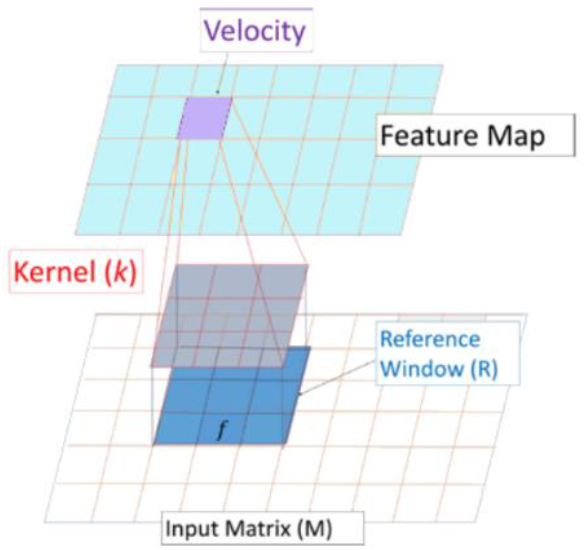
Determining the reference window for feature evaluation using a CNN kernel and kernel map.

We first calculated all the kernel maps using the weights from the first convolutional layer for each sample in the test set which was classified correctly. Next, we looked at the feature maps generated by a single kernel, *k*, across all the samples for a specific class, c. For each of these feature maps, we took the top 10% of the maximum values. For each velocity selected, we traced its location in the feature map *(u, v)* back to the submatrix of the input **M** from which it was calculated. We call this matrix **R** our reference window. More specifically, given a kernel *k* with weights *W*^(*k*)^ with dimensions *r × s*,

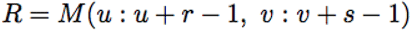

Within the reference window, every position (*i,j*) is equivalent to some node *v* from the phylogenetic tree with an taxon label, *f*. We calculate the importance of each feature *f* given the reference window **R** for sample *s* as its proportion of the velocity,

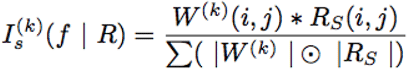

Here *k* is our current kernel with weights *W*^(*k*)^ which have been flipped to account for the convolution function; the summation is over all positions in **R**_s_. The absolute value of the weights in the denominator was used in order to handle any case where the contribution of one large positive component and one large negative component can give rise to a velocity that is much smaller than its components. This could lead to sporadic scaling of importance values which would be hard to interpret. By using the absolute value of the weights, we created an upper boundary of 1 and a lower boundary of −1. Within a single reference window, some taxa may have been found important in a small subset of the samples but may not be important considering all of the samples. In order to capture only the taxa which were consistently found important, we calculated the mean importance value of a feature f across all samples in class *c* given a single reference window **R** and kernel *k*.

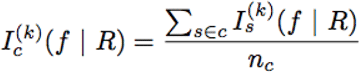

Since the reference windows of different velocities may overlap, it is possible for a single feature to have multiple importance values using the same kernel. A feature may also be found to be important by multiple kernels. This leads to multiple importance values for a single feature. To handle this problem, we selected the importance of f to be the maximum over all reference windows containing f and over all kernels, *k*.

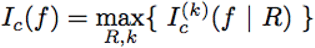

Lastly, we assign a score for a feature from the perspective of class *c* as the difference of the feature importance using all the samples within the class and the feature importance using all the samples not in the class. Given only two classes, the scores will be the same values with opposite sign. Despite that, we designed our method to be able to handle scenarios where there are more than two classes.

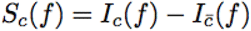

From these scores we created a list of feature scores for each class, allowing the analysis of feature importance from the perspective of different classes that can then be ranked. If a taxon was not found in any of the kernels, it is ranked at the bottom the feature list. To create a single ranked list, we then ranked features by the maximum score across classes.

### Model Prediction and Feature Aggregation

Models were trained using 10-fold cross-validation and report the average and standard deviation of each evaluating criterion from 10 times cross-validations. In each 10-fold cross-validation, the dataset was partitioned into 10 sets stratified by class proportion. We then constructed 10 datasets where a single partition was left out as the test set and the remaining were used for the training set.

The prediction evaluation for each dataset using the original taxa is shown in Table S1.

**Table S1.** Evaluation of each model on the five datasets using 10 iterations of 10-fold cross-validation.

|  |  | RF | SVM | LASSO | MLPNN |
| --- | --- | --- | --- | --- | --- |
| Cirrhosis | AUC | **0.944 (0.057)** | 0.815 (0.085) | 0.916 (0.071) | 0.918 (0.070) |
|  | MCC | **0.762 (0.021)** | 0.612 (0.061) | 0.527 (0.063) | 0.703 (0.137) |
|  | Precision | **0.881 (0.058)** | 0.763 (0.097) | 0.826 (0.095) | 0.843 (0.073) |
|  | Recall | **0.871 (0.061)** | 0.749 (0.095) | 0.814 (0.094) | 0.824 (0.074) |
|  | F1 | **0.870 (0.061)** | 0.745 (0.097) | 0.812 (0.095) | 0.822 (0.075) |
| T2D | AUC | **0.738 (0.060)** | 0.599 (0.071) | 0.646 (0.071) | 0.692 (0.059) |
|  | MCC | **0.334 (0.110)** | 0.141 (0.140) | 0.194 (0.125) | 0.278 (0.121) |
|  | Precision | **0.669 (0.056)** | 0.572 (0.071) | 0.598 (0.063) | 0.641 (0.061) |
|  | Recall | **0.665 (0.054)** | 0.569 (0.069) | 0.596 (0.061) | 0.637 (0.059) |
|  | F1 | **0.663 (0.054)** | 0.567 (0.070) | 0.594 (0.062) | 0.635 (0.059) |
| Obesity | AUC | **0.657 (0.123)** | 0.600 (0.119) | 0.575 (0.092) | 0.650 (0.104) |
|  | MCC | 0.056 (0.192) | 0.035 (0.162) | 0.022 (0.190) | **0.201 (0.178)** |
|  | Precision | 0.554 (0.154) | 0.532 (0.140) | 0.535 (0.133) | **0.638 (0.083)** |
|  | Recall | 0.647 (0.047) | **0.635 (0.052)** | 0.627 (0.056) | 0.631 (0.080) |
|  | F1 | 0.549 (0.066) | 0.540 (0.056) | 0.547 (0.071) | **0.629 (0.079)** |
| IBD | AUC | **0.899 (0.103)** | 0.859 (0.130) | 0.817 (0.151) | 0.893 (0.116) |
|  | MCC | 0.307 (0.321) | 0.417 (0.355) | 0.215 (0.332) | **0.552 (0.316)** |
|  | Precision | 0.754 (0.144) | 0.799 (0.137) | 0.713 (0.140) | **0.849 (0.118)** |
|  | Recall | 0.818 (0.065) | 0.818 (0.100) | 0.792 (0.082) | **0.835 (0.109)** |
|  | F1 | 0.765 (0.095) | 0.797 (0.113) | 0.739 (0.101) | **0.832 (0.111)** |
| CRC | AUC | - | - | - | - |
|  | MCC | **0.339 (0.197)** | 0.315 (0.174) | 0.242 (0.192) | 0.261 (0.195) |
|  | Precision | 0.534 (0.154) | **0.540 (0.143)** | 0.505 (0.207) | 0.524 (0.146) |
|  | Recall | **0.553 (0.125)** | 0.539 (0.110) | 0.468 (0.107) | 0.504 (0.127) |
|  | F1 | **0.522 (0.129)** | 0.513 (0.118) | 0.404 (0.136) | 0.492 (0.130) |

In our experiments, we observed that RF models performed the best overall compared to the other models. However, within individual partitions of a 10 times 10-fold cross validation evaluation, we observed that there were partitions in which RF models were outperformed by other models. An example of this for the obesity dataset is shown in Fig. S5. Because of this observation, we decide to use features from multiple learning models across multiple learning models. In this way, we believe that the true signal will be consistent between models and will be ranked higher than any signal from over-fitting when all the rankings are aggregated.

**Figure S5.**
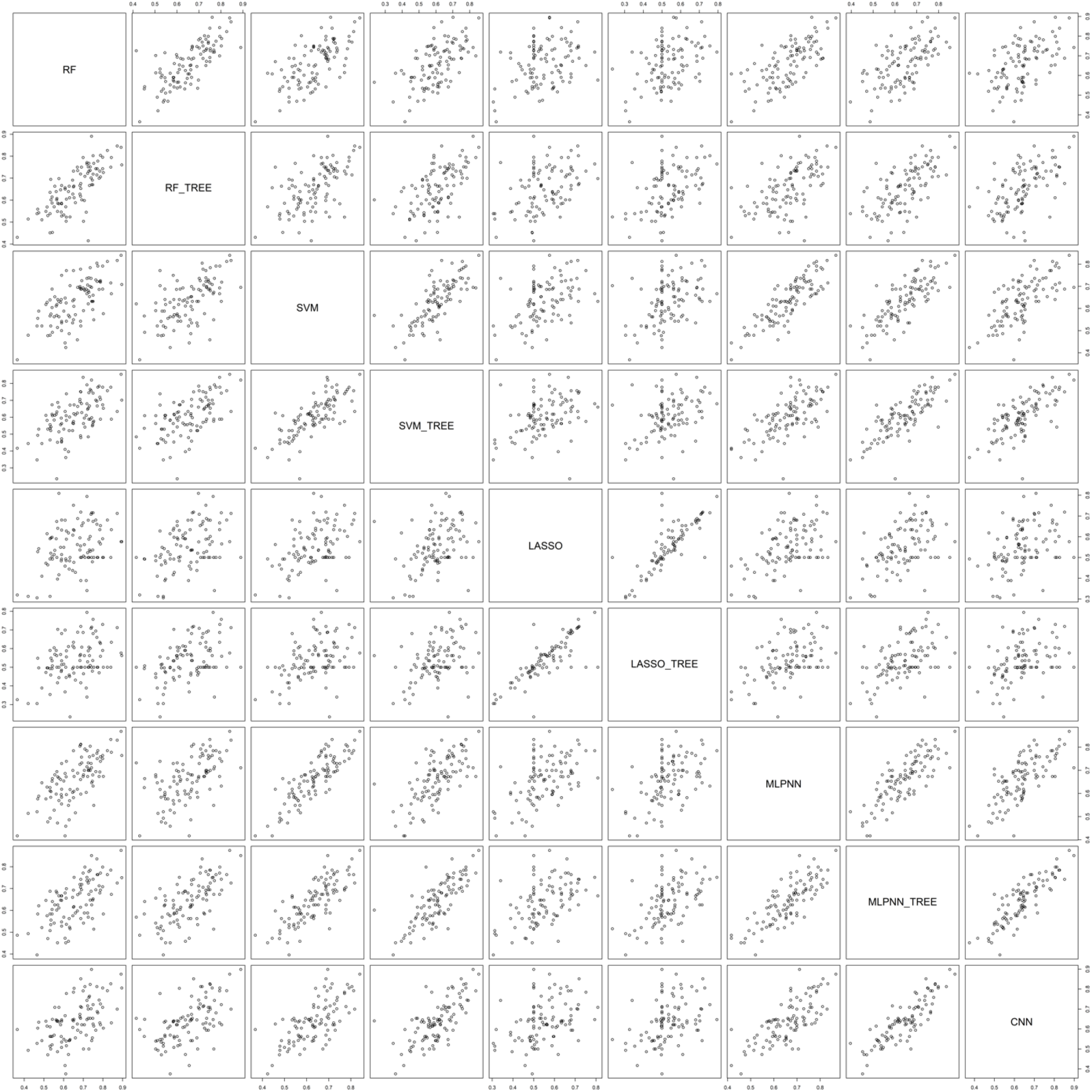
AUC scatterplots for pair-wise combinations of learning models on the obesity dataset.

### The original and tree features extracted from Meta-Signer

We report the aggregated top 20 original and tree-level taxa for each of the five datasets as well as the proportion of individual ranked lists in which a taxon was found in the top 20 features in Tables S2–S6.

**Table S2:**
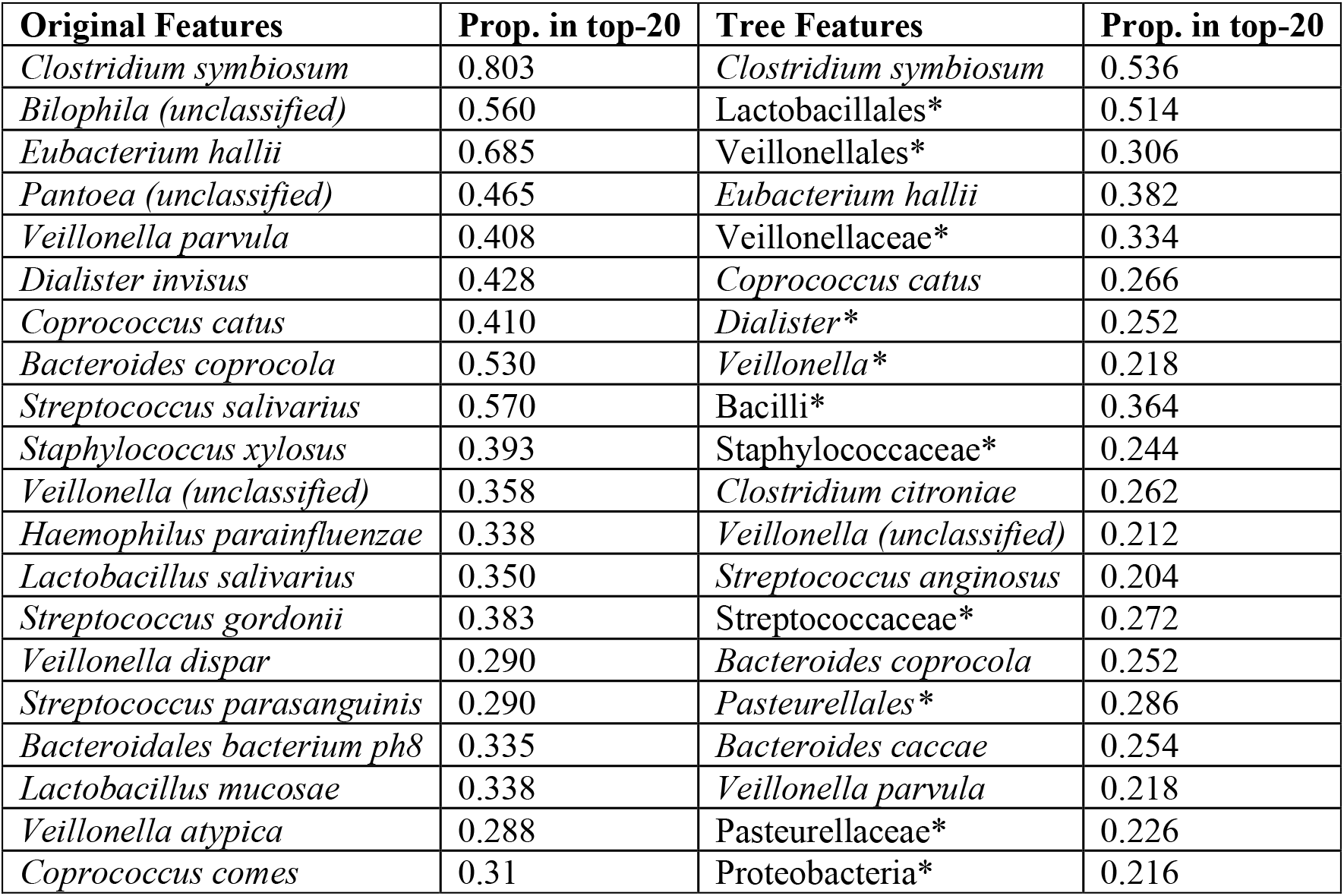
Cirrhosis original and tree features. Internal nodes on the tree are marked with an asterisk (*).

In the cirrhosis patients, four species of *Veillonella*, three species of *Streptococcus*, and *Haemophilus parainfluenzae* were found to be discriminative. In healthy patients, Meta-Signer identified *Eubacterium hallii*, two species of *Bacteroides* to be enriched in healthy patients. These findings agree with the original study. The tree taxa captured the Vellionellales order at the order level, family level, genus level, and species level. Meta-Signer also identified the tree branch from the class Bacilli to the genus *Streptococcus* as well as the species underneath. These two genera were identified as being strongly indicative of the disease state for cirrhosis in the original study.

**Table S3:**
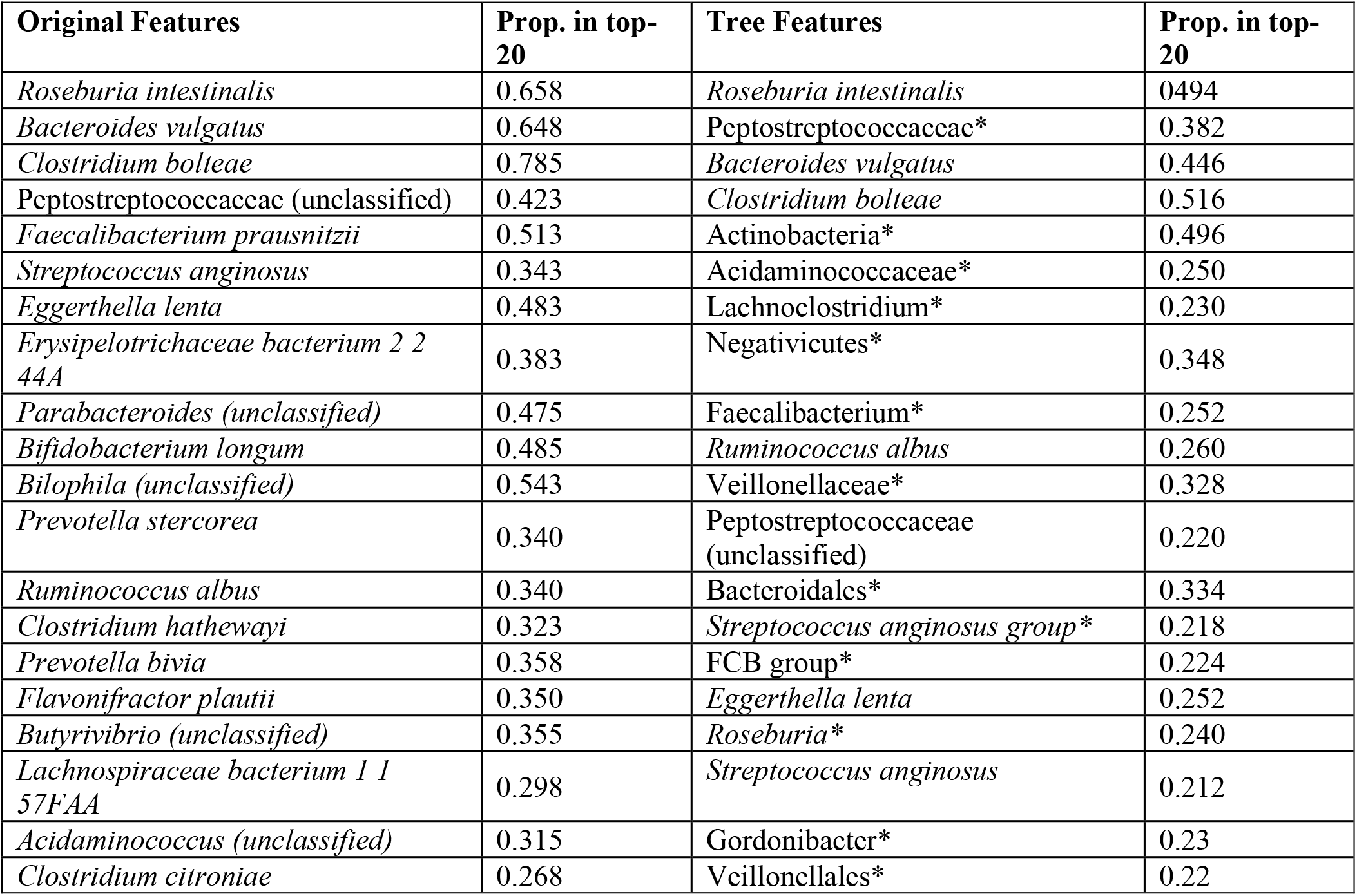
T2D original and tree features. Internal nodes on the tree are marked with an asterisk (*).

In patients with T2D, Meta-Signer identified *Roseburia intestinalis* and *Faecalibacterium prausnitzii* enriched in healthy patients while *Clostridium bolteae*, *Clostridium hathewayi*, and *Eggerthella lenta* were enriched in subjects with T2D. Similar observations were reported in previous studies. In addition, we observed an overall increase of the families Clostridaceae and Peptostretpcoccaceae in the tree. A previous study has reported that these two families of microbes are known to increase the blood levels of trimethylamine N-oxide (TMAO) in humans and that higher levels of TMAO are associated with diabetes.

**Table S4:** Obesity original and tree features. Internal nodes on the tree are marked with an asterisk (*).

| Original Features | Prop. in top-20 | Tree Features | Prop. in top-20 |
| --- | --- | --- | --- |
| *Deinococcus (unclassified)* | 0.775 | *Ruminococcus lactaris* | 0.682 |
| *Ruminococcus lactaris* | 0.700 | *Ruminococcus flavefaciens* | 0.494 |
| *Ruminococcus flavefaciens* | 0.615 | *Blautia producta* | 0.396 |
| *Alistipes senegalensis* | 0.54 | Deinococcaceae* | 0.380 |
| *Blautia producta* | 0.478 | *Alistipes senegalensis* | 0.468 |
| *Alistipes (unclassified)* | 0.478 | *Clostridium leptum* | 0.492 |
| *Ruminococcus torques* | 0.470 | *Ruminococcus bromii* | 0.332 |
| *Clostridium leptum* | 0.568 | *Alistipes (unclassified)* | 0.396 |
| *Ruminococcus bromii* | 0.450 | *Ruminococcus torques* | 0.392 |
| *Dorea formicigenerans* | 0.450 | Deinococcus-Thermus* | 0.312 |
| *Eubacterium siraeum* | 0.513 | *Dorea formicigenerans* | 0.306 |
| *Butyrivibrio crossotus* | 0.435 | Negativicutes* | 0.282 |
| *Mitsuokella (unclassified)* | 0.390 | *Butyrivibrio* * | 0.290 |
| *Bacteroides ovatus* | 0.348 | Deinococci* | 0.270 |
| *Bacteroides massiliensis* | 0.370 | *Bacteroides massiliensis* | 0.294 |
| *Akkermansia muciniphila* | 0.313 | *Mitsuokella (unclassified)* | 0.262 |
| *Veillonella (unclassified)* | 0.270 | Deinococcales* | 0.240 |
| *Desulfovibrio piger* | 0.325 | *Megasphaera elsdenii* | 0.224 |
| *Megasphaera elsdenii* | 0.298 | Micrococcaceae* | 0.214 |
| *Oscillibacter (unclassified)* | 0.253 | *Deinococcus (unclassified)* | 0.236 |

**Table S5:**
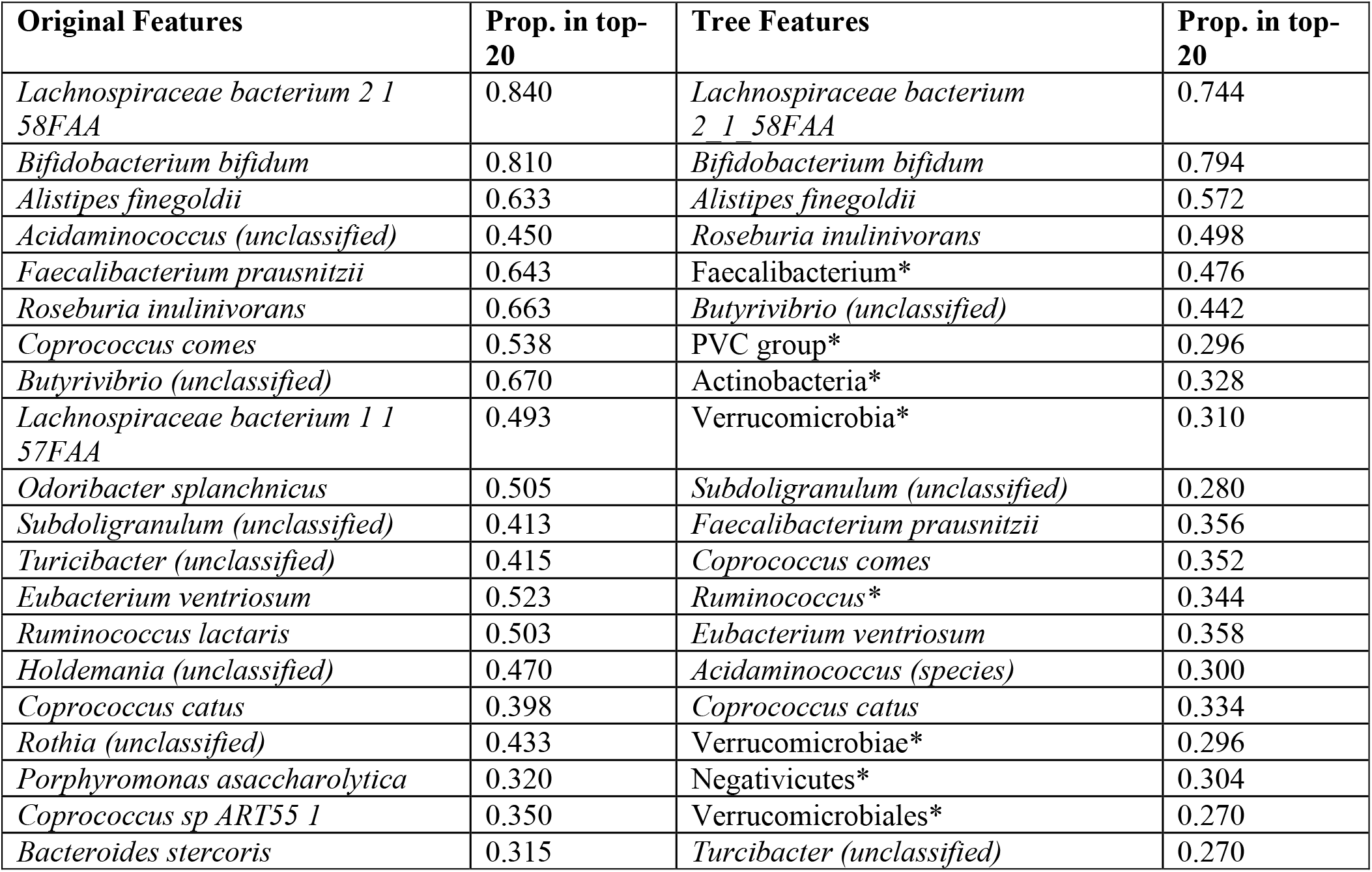
IBD original and tree features. Internal nodes on the tree are marked with an asterisk (*).

Meta-Signer finds microbes from the Lachnospiraceae family as well as *Bifidobacterium bifidum* and *Faecalibacterium prausnitzii* to be associated with patients with IBD. These findings agree with a separate analysis of this IBD dataset, however many studies have shown the same identified microbes as protective against IBD using different datasets.

**Table S6:** CRC original and tree features. Internal nodes on the tree are marked with an asterisk (*).

| Original Features | Prop. in top-20 | Tree Features | Prop. in top-20 |
| --- | --- | --- | --- |
| *Peptostreptococcus stomatis* | 0.685 | *Peptostreptococcus* * | 0.393 |
| *Parvimonas (unclassified)* | 0.600 | *Peptostreptococcus stomatis* | 0.397 |
| *Fusobacterium nucleatum* | 0.695 | *Tissierellia* * | 0.443 |
| *Parvimonas micra* | 0.590 | *Fusobacteria* * | 0.390 |
| *Eubacterium rectale* | 0.645 | *Fusobacteriales* * | 0.377 |
| *Ruminococcus bromii* | 0.590 | *Tissierellales* * | 0.397 |
| *Streptococcus salivarius* | 0.520 | *Eubacteriaceae* * | 0.543 |
| *Lachnospiraceae bacterium 5 1 63FAA* | 0.565 | *Peptoniphilaceae* * | 0.363 |
| *Gemella morbillorum* | 0.540 | *Parvimonas (unclassified)* | 0.367 |
| *Eubacterium hallii* | 0.515 | *Fusobacteriia* * | 0.400 |
| *Roseburia intestinalis* | 0.520 | *Terrabacteria group* * | 0.473 |
| *Bacteroides fragilis* | 0.495 | *Eubacterium* * | 0.383 |
| *Solobacterium moorei* | 0.485 | *Fusobacterium* * | 0.317 |
| *Lachnospiraceae bacterium 7 1 58FAA* | 0.450 | *Parvimonas* * | 0.313 |
| *Dorea formicigenerans* | 0.450 | *Burkholderiales* * (unclassified) | 0.323 |
| *Escherichia coli* | 0.400 | *Firmicutes* * | 0.327 |
| *Clostridium hathewayi* | 0.440 | *Fusobacteriaceae* * | 0.330 |
| *Anaerostipes hadrus* | 0.460 | *Lachnospiraceae* * | 0.287 |
| *Eubacterium eligens* | 0.260 | *Gammaproteobacteria* * | 0.223 |
| *Dorea longicatena* | 0.225 | *Lachnospiraceae (unclassified)* | 0.163 |

In patients with CRC, Meta-Signer found *Peptostreptococcus stomatis, Parvimonas micra, Fusobacterium nucleatum*, and *Streptococcus anginosus* to be enriched. All of these microbes have been associated with CRC in previous studies.

### Evaluation of the Meta-Signer features against other methods

We evaluated the extracted features from each method using 10-fold cross validation. Each dataset was randomly partitioned into 10 sets, stratified to balance the class proportion. Then each method was applied to 9 of the 10 sets, the training set, to extract features. The datasets were then filtered to only have the chosen features and models were trained on the training set and evaluated on the test. Each training set was used to train an SVM model over both the linear and Gaussian kernel. We observed that Meta-Signer was robust across all datasets. In cirrhosis, T2D, and obesity, it outperformed all other models. Biosigner and HFE were observed to perform very poorly for these datasets. Biosigner often returned no features (in which case an AUC of 0.5 was assigned to the testing). In the obesity dataset, both Biosigner and HFE extract features that lead to a testing AUC less than 0.5. We observed that a Wilcoxon test was the best for IBD and both Wilcoxon and HFE performed will in CRC. However, although not the best in IBD and CRC, Meta-Signer was still comparable. The table of cross-validated results across all learning methods and datasets is shown in Table S7.

**Table S7:** Meta-Signer performance including linear SVM features

|  | Meta-Signer | Meta-Signer<br>(Tree) | Biosigner | HFE | Wilcoxon |
| --- | --- | --- | --- | --- | --- |
| Cirrhosis | **0.903 (0.061)** | 0.871 (0.080) | 0.815 (0.231) | 0.627 (0.349) | 0.822 (0.168) |
| T2D | 0.647 (0.069) | **0.651 (0.065)** | 0.578 (0.052) | 0.584 (0.098) | 0.563 (0.138) |
| Obesity | **0.582 (0.134)** | 0.536 (0.146) | 0.437 (0.103) | 0.485 (0.127) | **0.582 (0.189)** |
| IBD | 0.782 (0.143) | 0.750 (0.189) | 0.639 (0.111) | 0.831 (0.091) | **0.875 (0.100)** |
| CRC | 0.332 (0.204) | 0.305 (0.193) | - | **0.339 (0.197)** | 0.246 (0.208) |

We then observed the extracted taxa from Meta-Signer and looked at the proportion of how often they appeared in the top 20 features in each model independently. We observed a large consensus in many taxa, however, a few taxa were missed by some models completely and picked up by the others. The top 20 taxa found in obesity using both raw and tree level features are shown in Tables S8 and S9.

**Table S8.** The top 20 original taxa found in the Cirrhosis dataset and the proportion of models that contained that feature in the top 20 of their own ranked list.

|  | RF | SVM | LASSO | MLPNN |
| --- | --- | --- | --- | --- |
| *Deinococcus (unclassified)* | 1 | 0.57 | 0.98 | 0.55 |
| *Ruminococcus lactaris* | 0.99 | 0.47 | 1 | 0.34 |
| *Ruminococcus flavefaciens* | 1 | 0.51 | 0.58 | 0.37 |
| *Alistipes senegalensis* | 1 | 0.05 | 0.72 | 0.39 |
| *Blautia producta* | 0 | 0.54 | 0.9 | 0.47 |
| *Alistipes (unclassified)* | 0.44 | 0.06 | 0.91 | 0.5 |
| *Ruminococcus torques* | 0.02 | 0.94 | 0.67 | 0.25 |
| *Clostridium leptum* | 0.69 | 0.63 | 0.56 | 0.39 |
| *Ruminococcus bromii* | 0.97 | 0.11 | 0.43 | 0.29 |
| *Dorea formicigenerans* | 0.6 | 0.03 | 0.91 | 0.26 |
| *Eubacterium siraeum* | 0.88 | 0.5 | 0.44 | 0.23 |
| *Butyrivibrio crossotus* | 0.31 | 0.35 | 0.77 | 0.31 |
| *Mitsuokella (unclassified)* | 0 | 0.72 | 0.53 | 0.31 |
| *Bacteroides ovatus* | 0.93 | 0.14 | 0.1 | 0.22 |
| *Bacteroides massiliensis* | 0.62 | 0 | 0.62 | 0.24 |
| *Akkermansia muciniphila* | 0.86 | 0 | 0.25 | 0.14 |
| *Veillonella (unclassified)* | 0.99 | 0.02 | 0 | 0.07 |
| *Desulfovibrio piger* | 0.01 | 0.73 | 0.34 | 0.22 |
| *Megasphaera elsdenii* | 0 | 0.11 | 0.7 | 0.38 |
| *Oscillibacter (unclassified)* | 0.22 | 0.69 | 0.04 | 0.06 |

**Table S9.** The top 20 tree-level taxa found in the obesity dataset and the proportion of models that contained that feature in the top 20 of their own ranked list.

|  | **RF** | **SVM** | **LASSO** | **MLPNN** | **CNN** |
| --- | --- | --- | --- | --- | --- |
| *Ruminococcus lactaris* | 0.89 | 0.87 | 0.98 | 0.43 | 0.24 |
| *Ruminococcus flavefaciens* | 0.96 | 0.72 | 0.47 | 0.25 | 0.07 |
| *Blautia producta* | 0.01 | 0.61 | 0.96 | 0.4 | 0 |
| Deinococcaceae | 0.71 | 0 | 0.99 | 0.08 | 0.12 |
| *Alistipes senegalensis* | 0.87 | 0.28 | 0.77 | 0.33 | 0.09 |
| *Clostridium leptum* | 0.41 | 0.92 | 0.64 | 0.34 | 0.15 |
| *Ruminococcus bromii* | 0.89 | 0 | 0.41 | 0.34 | 0.02 |
| *Alistipes (unclassified)* | 0.28 | 0.41 | 0.92 | 0.35 | 0.02 |
| *Ruminococcus torques* | 0 | 0.85 | 0.72 | 0.2 | 0.19 |
| Deinococcus-Thermus | 0.79 | 0 | 0.01 | 0.23 | 0.53 |
| *Dorea formicigenerans* | 0.23 | 0.01 | 0.87 | 0.38 | 0.04 |
| Negativicutes | 0.17 | 0 | 0.46 | 0.08 | 0.7 |
| *Butyrivibrio* | 0.73 | 0 | 0.61 | 0.11 | 0 |
| Deinococci | 0.74 | 0 | 0.03 | 0.21 | 0.37 |
| Bacteroides massiliensis | 0.43 | 0.05 | 0.71 | 0.28 | 0 |
| *Mitsuokella (unclassified)* | 0 | 0.61 | 0.49 | 0.21 | 0 |
| Deinococcales | 0.75 | 0 | 0.04 | 0.12 | 0.29 |
| *Megasphaera elsdenii* | 0 | 0.12 | 0.66 | 0.34 | 0 |
| Micrococcaceae | 0.42 | 0 | 0.49 | 0.12 | 0.04 |
| *Deinococcus (unclassified)* | 0.71 | 0 | 0.1 | 0.2 | 0.17 |

We observed that the CNN model was not contributing as much to the top 20 selected features in each dataset. Upon visual inspection, we observed that the features from the CNN models were more often higher taxonomic levels (class and family levels and super levels). Therefore, we suspected that it captured information at higher aggregated points, and as such, less features are required. Therefore, we trained SVM models using just the top 10 features after aggregating the ranked lists from the CNN models only and compared the performance to Meta-Signer on the tree features. With the exception of the IBD dataset, the features extracted just from the CNN are comparable or even superior to the ensemble aggregated features. This makes us believe that the CNN is finding useful features, however the features it finds are being dampened by the other models in the ensemble aggregation. A table comparing the top 10 features of the CNN model and Meta-Signer is shown in Table S10.

**Table S10.** AUC values for SVM models trained on the top 10 tree level features using Meta-Signer and just CNN rankings.

| **Dataset** | **Meta-Signer (Tree)** | **PopPhy-CNN** |
| --- | --- | --- |
| Cirrhosis | 0.721 (0.303) | 0.859 (0.072) |
| T2D | 0.645 (0.076) | 0.567 (0.109) |
| Obesity | 0.500 (0.090) | 0.592 (0.129) |
| IBD | 0.728 (0.170) | 0.674 (0.177) |
| CRC | 0.186 (0.221) | 0.143 (0.221) |

